# Prenatal corticosteroid exposure disrupts vascular-immune interactions and impairs steroidogenesis in the fetal testis

**DOI:** 10.64898/2026.01.27.702066

**Authors:** Satoko Matsuyama, Lauren Hudepohl, Kazuhiro Matsuyama, Shu-Yun Li, Meghana Ginugu, Xiaowei Gu, Matthew J. Kofron, Vikram Ravindra, Tetsuo Shoda, Tony DeFalco

## Abstract

Antenatal corticosteroid (ACS) therapy, typically using dexamethasone (Dex), is a cornerstone of improving preterm infant survival by accelerating lung maturation. However, its systemic effects on other developing organs remain poorly understood. Here, we reveal a previously unrecognized impact of prenatal Dex exposure on fetal testis development, with implications for long-term male reproductive health. Analysis of fetal testes from a Dex-based ACS mouse model revealed that Dex suppressed androgen synthesis by reducing Leydig cell number and downregulating steroidogenic pathways. Dex also activated immune programs, inducing CD163⁺ M2 macrophages and IL10 signaling, which promoted endothelial expansion. Despite these pro-angiogenic signals, vascular architecture was disrupted: capillary density and pericyte coverage declined, while blood vessel diameter increased. Transcriptomic analyses revealed downregulation of androgen response and cholesterol metabolism pathways, alongside upregulation of immune and coagulation signatures. Analyses of fetal ovaries revealed a sexually dimorphic and organ-specific effect of ACS therapy, in which ovarian gene expression remained unaffected. Our findings uncover an immune-vascular mechanism linking prenatal Dex exposure to impaired steroidogenesis, providing new insight into potentially broad effects of ACS therapy.

**One Sentence Summary:** Prenatal corticosteroid exposure disturbs immune-vascular interactions and suppresses androgen synthesis in the fetal testis, revealing potentially widespread effects of a commonly used therapy.

## Introduction

One of the major causes of early neonatal death among preterm infants is respiratory complications such as respiratory distress syndrome (RDS), which arise from structurally and functionally immature lungs. Complications of prematurity are a leading cause of both neonatal and under-5 mortality worldwide, and it is estimated that in 2020 approximately 13.4 million infants were born preterm, which represents about 10% of all live births (1). Improving survival and reducing the morbidity of preterm infants require not only optimal postnatal intensive care, but also effective promotion of fetal lung maturation in utero, which remains a global challenge. To this end, antenatal corticosteroid (ACS) administration has been widely implemented as a standard of care for women at high risk of preterm birth in many countries, particularly high-income settings including the United States, where >90% of women at risk of delivery between 24- and 34-weeks’ gestation receive some form of ACS, most commonly betamethasone or dexamethasone. Dexamethasone (Dex), a synthetic glucocorticoid (GC), is one of the prototypical agents used as an ACS and has been shown to accelerate fetal lung maturation, thus significantly reducing the incidence of RDS and neonatal death. These benefits have been established since the seminal trial by Liggins and colleagues and confirmed in numerous subsequent randomized controlled trials and Cochrane reviews (2, 3). However, the long-term safety of ACS, including its effects on non-pulmonary organs and neurodevelopment, as well as its potential impact on infants who ultimately deliver in the late preterm period or at term despite having been judged at risk and exposed to ACS, remains incompletely understood. It has also been suggested that this treatment may modulate the immune system, alter glucose sensitivity, and delay thymic development, but these effects have not been fully elucidated (4).

Glucocorticoids regulate multiple essential systemic functions, including blood pressure and immune responses. In mice, corticosterone is the predominant endogenous glucocorticoid (5), whereas in humans it is cortisol (6). Major downstream signaling pathways are mediated by the glucocorticoid receptor (GR), a member of the nuclear receptor superfamily whose official name is NR3C1. Upon binding to cortisol, GR translocates to the nucleus, where it interacts with specific DNA elements to regulate gene transcription (7). GR function is further fine-tuned by mechanisms that control its intracellular trafficking, promoter specificity, cofactor interactions, receptor stability, and turnover (8). Cellular responses to glucocorticoids are governed by the recruitment of cofactors that act through diverse mechanisms, including chromatin remodeling, assembly of the transcriptional machinery, and post-translational modification of histones and other components of transcription factor complexes (9–11). Tissue-specific GR knockout models have been generated in multiple organs (12), but studies employing testis-specific GR knockout models remain scarce, and, thus, the role of GR in the testis is still poorly defined.

Although GR expression and cellular localization in the postnatal and adult testis have been described, its functional roles have been addressed in only a limited number of cell-type-specific perturbation studies. In particular, very little is known about its expression pattern and role during prenatal testis development. In postnatal day 20 mouse testes, GR expression has been reported in Leydig cells, peritubular myoid cells, Sertoli cells, and early germ cells (13). In adult mice, GR is expressed in the nuclei of spermatogonia and preleptotene spermatocytes, whereas its expression is weaker in pachytene spermatocytes and virtually absent in spermatids (14). Sertoli-cell-specific deletion of *Gr* in mice has little to no overt impact on fertility yet reduces the number of Sertoli cells and spermatocytes (13). In addition, a ∼50% reduction of GR in Leydig cells achieved by AAV9-Cre-mediated targeting impairs steroidogenesis (15), suggesting that GR is required in adult Leydig cells. Nevertheless, GR localization and function during fetal stages of testis development remain largely unexplored.

Excessive exposure to glucocorticoids can also have profound effects on male reproductive development. Synthetic glucocorticoids such as betamethasone and dexamethasone disrupt male reproductive hormones across species. In medaka, betamethasone treatment induces feminization (16). In mice, dexamethasone administration during critical windows of embryonic development increases the expression of male-specific genes such as *Sry* and *Sox9* (17). In rats treated with glucocorticoids during prenatal development, nuclear COUP-TFII (NR2F2), a marker of Leydig progenitor cells, is upregulated, while proliferation of stem Leydig cells is reduced and testosterone levels are decreased (18). Prenatal dexamethasone exposure in rats has been shown to impair adult testicular structure and function. In one study, dexamethasone was administered from gestational day 9 to 20, and testes were examined in adulthood: testicular volume and weight were reduced, the interstitial area between seminiferous tubules was expanded, and sperm counts were decreased (19). Epigenetic reprogramming affecting sperm quality has also been reported (20). Collectively, these observations highlight the need for further investigation into the long-term consequences of fetal glucocorticoid therapy on male reproductive health.

Here we show that the response to prenatal dexamethasone differs between testes and ovaries, with no detectable effect in ovaries. In the fetal testis, prenatal dexamethasone administration activated the immune system, lowered androgen levels, and disrupted vascular patterning. Mechanistically, dexamethasone drove M2 macrophages toward an M2c phenotype characterized by induction of CD163. M2c macrophages are induced in the presence of interleukin 10 (IL10), transforming growth factor beta (TGF-β), and glucocorticoids (21, 22). They are usually considered immunoregulatory (anti-inflammatory) macrophages and involved in phagocytosis of apoptotic cells (21, 22). M2c macrophages secrete large amounts of IL10 and TGF-β, and express multiple markers including CD163, CD206, RAGE, and other scavenger receptors (21). Consistent with reports that CD163+ macrophages promote angiogenesis and vascular function (23), we show that CD163-positive macrophages promote angiogenesis, and that macrophage-derived IL10 further supports angiogenic remodeling; ex vivo gonad culture experiments confirmed a pro-angiogenic role for IL10 in fetal testes. Prenatal dexamethasone also increased VCAM1 expression and enhanced monocyte recruitment. Despite these pro-angiogenic signals, angiogenesis was ultimately disrupted: capillary density and pericyte coverage were reduced, leading to vessel dilation and a higher proportion of large-diameter vessels. This altered vascular architecture was associated with a decrease in Leydig cell number and function as vascular area increased, suggesting that prenatal dexamethasone-induced vascular remodeling contributes to impairment of testicular steroidogenesis.

## Results

### Glucocorticoid receptor (GR) is widely expressed during fetal testis development

To determine the localization of GR during fetal testis development, we examined GR expression in various cell types. At embryonic day 12.5 (E12.5), GR was absent in the testis (Fig. S1A). By E13.5, GR remained undetectable in the testis but was expressed in the neighboring mesonephros (Fig. S1A). From E14.5 onward, GR was expressed in Leydig cells, interstitial progenitors, germ cells, endothelial cells, macrophages, and monocytes (Fig. 1A, 1B). Macrophages were defined as F4/80⁺CD45⁺ cells, and monocytes as F4/80⁻CD45⁺ cells (24). Sertoli cells did not express GR during fetal stages (Fig. 1A, 1B). At E16.5, GR expression was expressed in an increasing number of interstitial progenitors, germ cells, and endothelial cells (Fig. 1B). Germ cells exhibited variable GR expression, with either strong or weak expression (Fig. 1C). Thus, GR is widely expressed in the late fetal testis.

**Figure 1.**
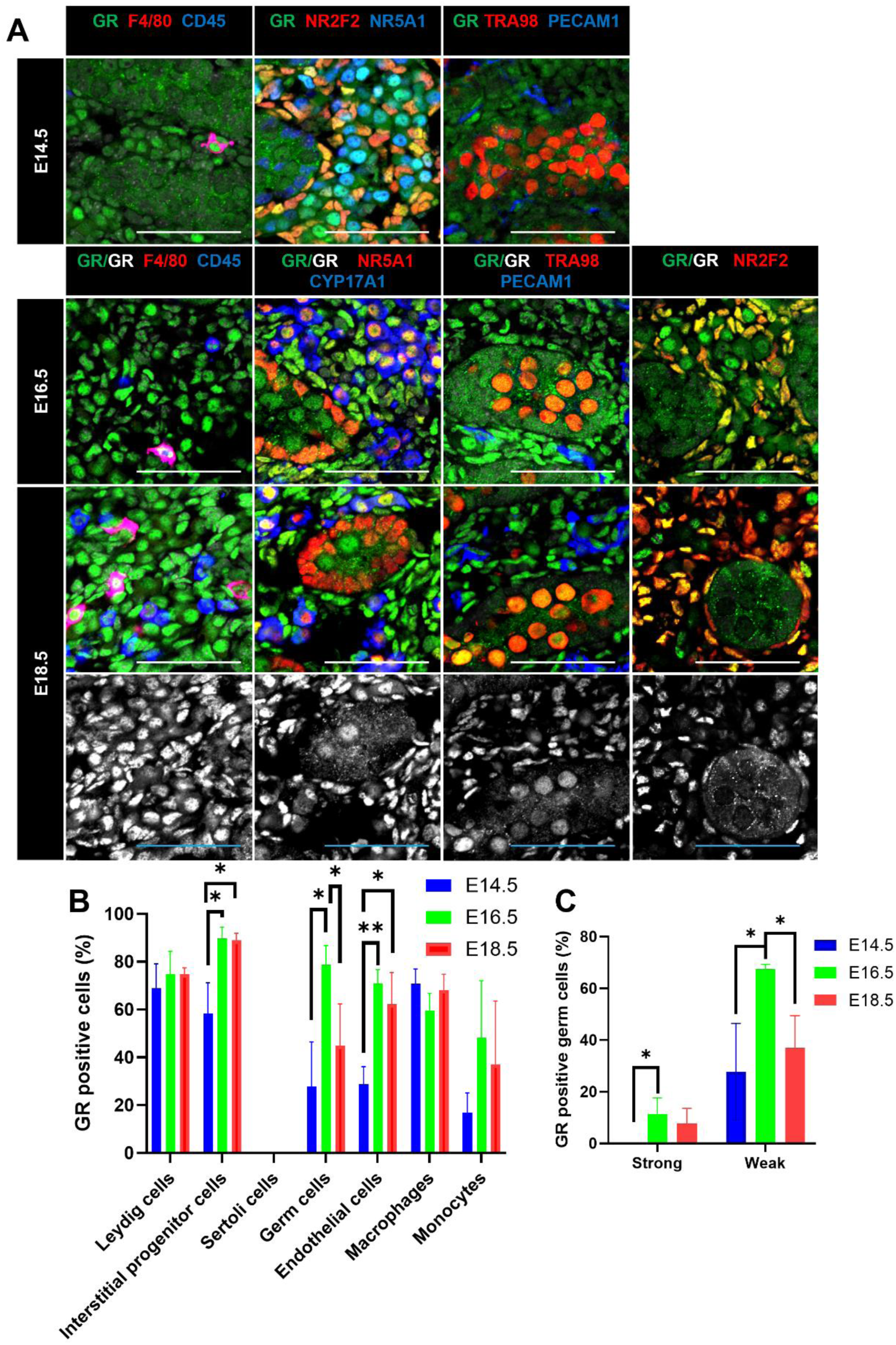
GR is widely expressed during fetal testis development. (A) Immunofluorescence images of E14.5, E16.5, and E18.5 testes showing GR localization with cell-type-specific markers (NR5A1: Sertoli and interstitial mesenchymal cells; TRA98: germ cells; CYP17A1: Leydig cells; NR2F2: Leydig progenitors; PECAM1: germ and endothelial cells; F4/80: macrophages; CD45: pan-immune cells). (B) Quantification of GR-positive Leydig cells, interstitial progenitors, Sertoli cells, germ cells, endothelial cells, macrophages, and monocytes at E14.5, E16.5, and E18.5. (C) Quantification of percent germ cells with strong versus weak GR expression. Scale bar: 100 μm. \**P*<0.05; \*\**P*<0.01.

### Prenatal dexamethasone activates testicular immune cells and suppresses androgen production

To elucidate the effects of late-term dexamethasone (Dex) exposure on the gonads, we developed a mouse model for fetal Dex therapy. In mice, Dex at 2-4 mg/kg/day represents a typical high-dose preclinical range (25, 26). Furthermore, due to the fast metabolic rate of mice, our 4 mg/kg/day dose yields an area under the plasma concentration-time curve (AUC) comparable to a standard human prenatal corticosteroid administration course. Saline, Dex 2 mg/kg/day, or Dex 4 mg/kg/day were administered to C57BL/6J (B6) pregnant mice from E14.5 to E17.5, and testes were collected at E18.5 (Fig. 2A). Compared to saline-treated controls, the number of NR5A1-positive and CYP17A1-positive cells (Leydig cells) was significantly reduced in both Dex 2 mg/kg/day and Dex 4 mg/kg/day groups (Fig. 2C). CYP17A1 expression was further decreased in the Dex 4 mg/kg/day group (Fig. 2B, 2F). Expression of CYP11A1 and STAR was also reduced (Fig. S2A, S2B), while Caspase 3 levels remained unchanged (Fig. S3B). Therefore, prenatal Dex suppresses androgen production. Furthermore, Dex 4 mg/kg/day treatment increased the number of CD45⁺ cells (Fig. 2D, 2E). Macrophage (F4/80^+^) numbers remained stable, whereas monocytes (F4/80^-^) increased significantly (Fig. 2E). Ki67⁺ cell numbers did not change (Fig. S3C). Quantitative real-time PCR (qRT-PCR) analysis confirmed significant upregulation of *Ptprc* (encoding CD45) and *Itgam* (encoding CD11b, a myeloid immune cell marker) following Dex 4 mg/kg/day treatment (Fig. 2G). Therefore, prenatal Dex increases immune cell numbers. Other cell types were not affected by Dex treatment, as the numbers of interstitial progenitors, germ cells, and endothelial cells were unaffected (Fig. S2C). GR⁺ cell proportions remained unchanged across cell types (Fig. S2D), although the number of weakly GR-expressing germ cells increased significantly (Fig. S2E).

**Figure 2.**
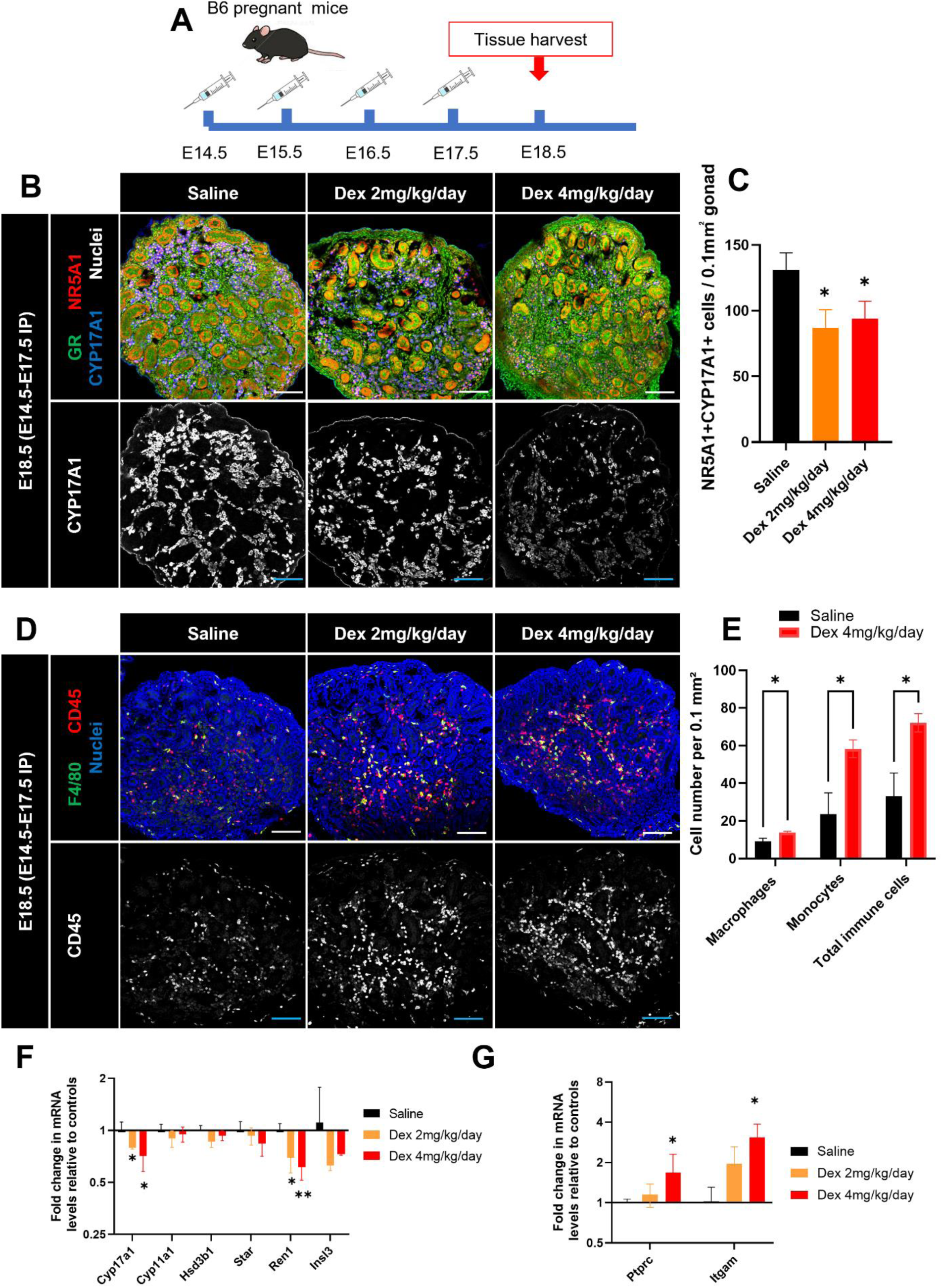
Prenatal dexamethasone reduces Leydig cell numbers and increases testicular immune cell number. (A) Cartoon of experimental design: saline, Dex 2 mg/kg/day, or Dex 4 mg/kg/day were administered daily from E14.5-E17.5, and testes were analyzed at E18.5. (B) Immunostaining for NR5A1 and CYP17A1. (C) Quantification of NR5A1⁺CYP17A1⁺ Leydig cells per 0.1 mm² gonadal area. (D) CD45 staining of immune cells. (E) Quantification of macrophages and monocytes per unit area. (F, G) qRT-PCR analysis of steroidogenic and immune-related genes. Scale bar: 100 μm. \**P*<0.05; \*\**P*<0.01.

### Dex treatment disrupts testicular vascular patterning

Analysis of PECAM1^+^ (CD31^+^) vascular endothelial cells revealed that Dex treatment reduced the number of thin blood vessels, resulting in a predominance of thicker vessels (Fig. 3A). ETS-related gene (ERG)-positive cells (endothelial cell nuclear marker) became sparse and showed altered spatial distribution, consistent with three-dimensional (3D) imaging results (Fig. S3A, Fig. 3B).

**Figure 3.**
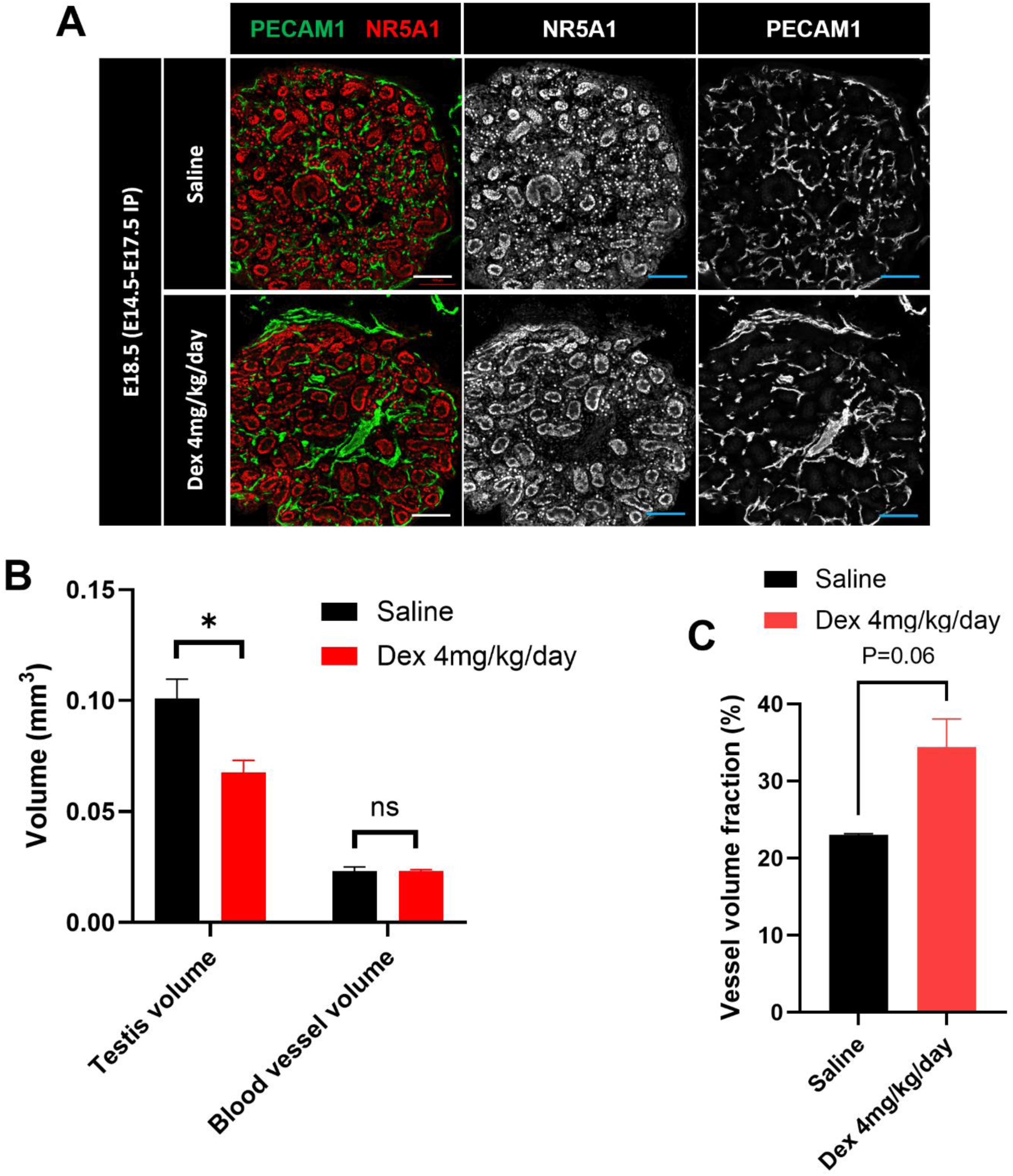
Glucocorticoid therapy disrupts fetal testicular vascular patterning. (A) PECAM1 staining of testes showing vascular architecture after saline or Dex treatment. (B) 3D tissue imaging and volumetric analyses of testis and vascular volume. (C) Quantification of percent vessel volume within the testis. Scale bar: 100 μm. \**P*<0.05.

Whole-mount 3D morphometric analyses using immunofluorescence revealed that testicular volume significantly decreased after Dex administration (Fig. 3B). Total vascular volume remained unchanged (Fig. 3B); therefore, the proportion of blood vessels within the testis increased (Fig. 3C). These results suggest that Dex disrupts vascular patterning in the fetal testis.

### Transcriptomic analysis reveals testicular immune activation and androgen suppression

To broadly examine the effects of Dex exposure, we performed bulk RNA-Seq of E18.5 fetal gonads from B6 pregnant dams injected with saline or 4 mg/kg/day Dex daily from E14.5 to E17.5. Samples were collected from three different dams treated either with saline or Dex, with each sample comprising four testes or eight ovaries. Testes from Dex-treated fetuses showed distinct transcriptomic profiles compared to controls, with 262 differentially expressed genes (DEGs) identified (Fig. 4A). A PCA plot confirmed clear group separation between Dex and control groups (Fig. S4A). Among the top 20 upregulated DEGs, five genes—*Slpi*, *Trem1*, *S100a8*, *S100a9*, and *Il36g*—were associated with macrophages and monocytes (Fig. S4B). Immune-related genes *Itgam, Trem1, Csf3r, S100a8*, and *S100a9* were upregulated. Among vascular-related genes, *Rgs5* and *Angpt2* were downregulated, while *Vcam1* and *Mmp9* were upregulated. Among Leydig cell genes, *Hmgcr, Hmgs1*, and *Hsd17b7* were downregulated, while *Dik1* was upregulated (Fig. 4A).

**Figure 4.**
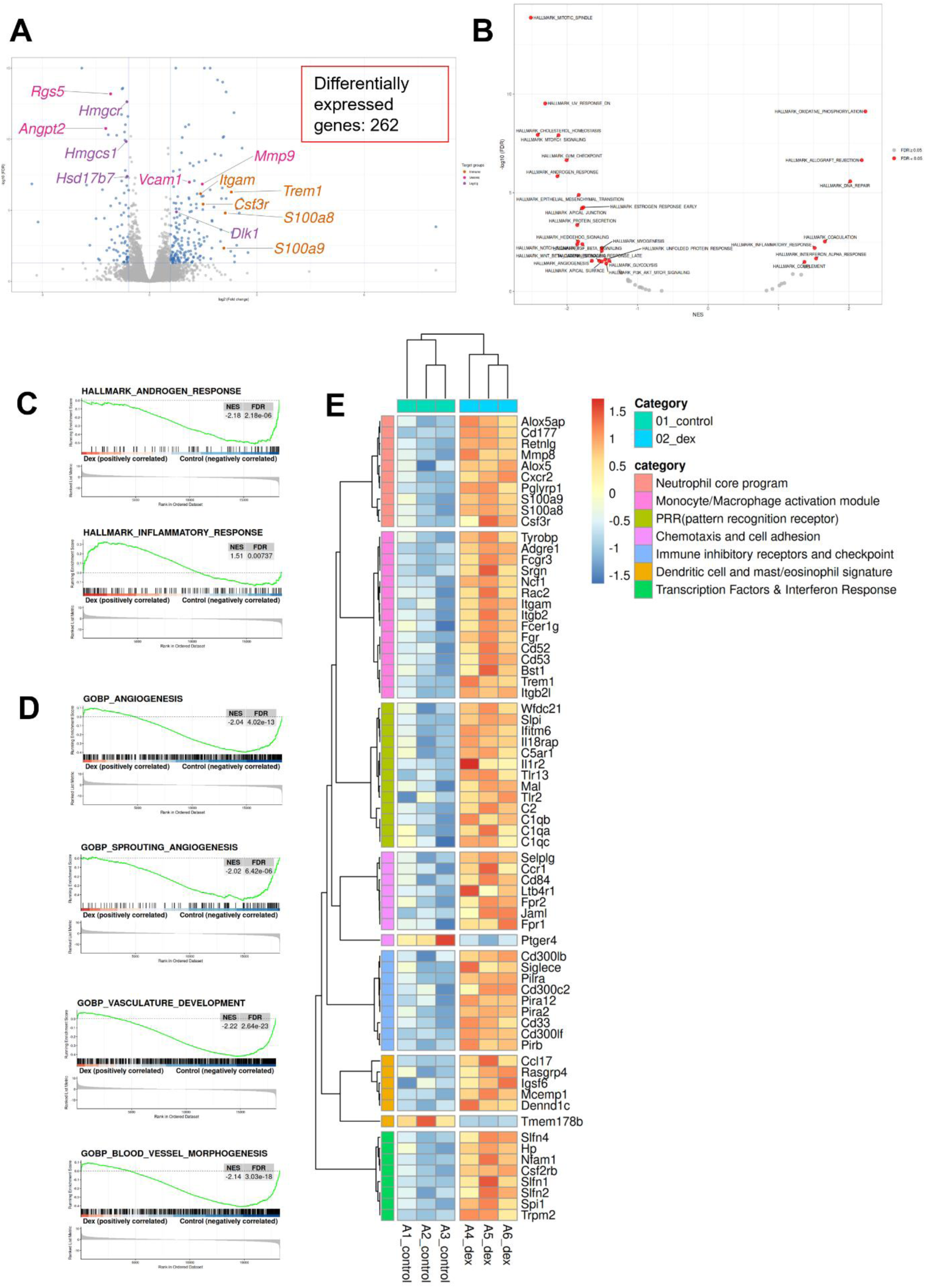
Transcriptomic analysis reveals immune activation and androgen suppression after Dex treatment. (A) Volcano plot of differentially expressed genes from whole-gonad bulk RNA-seq of E18.5 Dex-treated versus saline-treated control testes. Genes within specific developmental pathways are highlighted in different colors. (B) Volcano plot of GSEA analysis. (C) GSEA analysis of androgen response and inflammatory response pathways. (D) GSEA analysis of vascular development pathways. (E) Heatmap showing genes activated by Dex treatment within various immune modules.

Gene Set Enrichment Analysis (GSEA) revealed significant downregulation of androgen response, cholesterol homeostasis, and PI3K-AKT-mTOR signaling, alongside upregulation of coagulation (Fig. 4B). Individual GSEA analyses confirmed decreased androgen response and increased inflammatory response (Fig. 4C). Vascular system analyses indicated reduced angiogenesis (Fig. 4D).

Dex upregulated immune cell gene expression broadly (Fig. 4E). First, chemotaxis and cell adhesion pathways, such as *Ccr1*, were activated. Next, pattern recognition receptors (PRRs), including *Il1r2*, were upregulated. The neutrophil core program, indicated by *S100a8* and *S100a9*, were also upregulated, followed by activation of the monocyte/macrophage module with increased expression of *Adgre1* (encoding F4/80) and *Itgam*. In addition, immune inhibitory receptors and checkpoints, dendritic cell and mast/eosinophil signatures, and transcription factor/interferon response programs were all enhanced.

Angiogenesis and endothelial gene signatures were consistent with an impact on vascular remodeling. *Vcam1* and other endothelial interface molecules were upregulated, creating a platform to potentially facilitate leukocyte adhesion and migration. Angiogenic regulators were also affected. Additionally, the pericyte marker *Rgs5* was downregulated, suggesting a reduction in pericyte coverage and, thus, leading to a broader distribution of vessel diameter. Taken together, the results suggest that Dex reduces sprouting and branching while promoting vessel enlargement (Fig. 5A).

**Figure 5.**
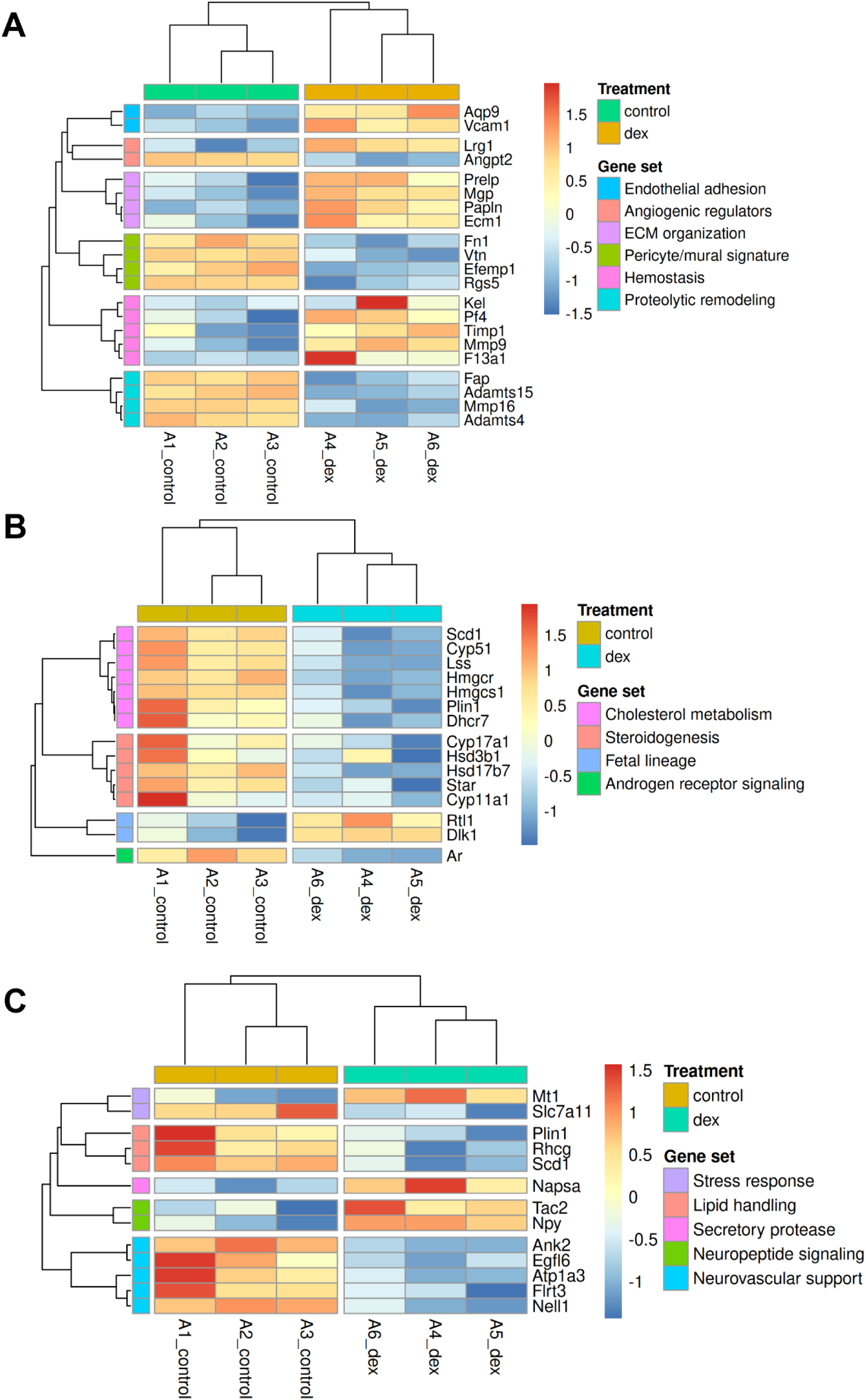
Dex reduces vascular sprouting and branching while promoting vessel enlargement and concurrently suppresses cholesterol metabolism and steroid production. (A) Heatmap of endothelial and angiogenic gene signatures. (B) Heatmap of cholesterol metabolism and steroidogenesis-associated genes impacted by Dex treatment. (C) Additional gene signature modules affected by Dex treatment.

Dex reduced cholesterol metabolism and steroid production, indicating Leydig dysfunction, since steroid production takes place in Leydig cells. Cholesterol metabolism-related genes and steroidogenesis-related genes were downregulated (Fig. 5B), suggesting that Dex reduces cholesterol metabolism and steroid production. Additional RNA-seq results are shown in Fig. 5C, S4, S5 and S6.

### Dex treatment has no effect on ovarian gene expression

In addition to the testis, we also assessed GR expression in the fetal ovary. In the ovary, GR was expressed in mesonephric tissue at E12.5, which increased at E13.5 and E14.5, reaching a peak at E14.5 (Fig. S7A, S7B). Thereafter, GR expression declined at E16.5 (Fig. S7B) and was generally weak through the entire ovary at E18.5 (Fig. S8A). Following Dex treatment, the number of endothelial cells increased (Fig. S8C), and the number of GR^+^ endothelial cells increased (Fig. S8D). Macrophages and monocytes in the ovaries showed few GR-positive cells, and no significant change was observed even after Dex treatment (Fig. S8B-D). RNA-seq analysis of ovaries revealed no significant differences between Dex-treated and control ovaries, and DESeq2-identified DEGs were minimal (Fig. S9A-E). Testes and ovaries were compared using specimens from the same dam, thus minimizing effects from variability due to different dams. To summarize, response to prenatal Dex differed between testes and ovaries, with no significant effect in ovaries.

### Dex treatment activates the IL10 pathway in testicular CD45⁺ cells

Whole-testis bulk RNA-seq analyses revealed significant alterations in immune-cell-related gene expression. Because immune cells constitute only a minor fraction of the total testicular cell population, we performed purification of E18.5 testicular CD45-positive cells for subsequent bulk RNA-seq analyses. This approach was designed to enrich not only macrophages and monocytes, but also other immune cell types present in the testis.

Following maternal administration of saline or Dex, CD45⁺ cells were sorted from fetal testes using magnetic-activated cell sorting (MACS) (Fig. 6A). Dex-treated samples separated clearly from controls in principal component analysis (Fig. S11A). Overall data quality and replicate concordance were supported by library distributions, MA-plot assessment, and sample-to-sample distance analyses (Fig. S10A-C). RNA-seq analyses of CD45-positive cells revealed 1125 differentially expressed genes between Dex-treated and control testes. GR-responsive genes such as *Tsc22d3* and *Fkbp5* were upregulated, demonstrating an intrinsic GR-driven response in immune cells. By sorting and specifically analyzing CD45^+^ cells, additional immunological changes became evident, in which immune cell gene expression significantly changed (Fig. 6B); in particular, *Cd163* and *Il10* were strongly upregulated (Fig. 6B). Transcriptome-wide differential expression was consistent across replicates, as shown by the hierarchical clustering heatmap (Fig. S11B). Canonical GR-responsive genes and immune-related gene sets exhibited coordinated regulation following Dex exposure (Fig. S11C). Additionally, the *Il10* pathway was activated in CD45⁺ cells following Dex treatment (Fig. 6C), indicating a shift towards an M2c-like macrophage phenotype.

**Figure 6.**
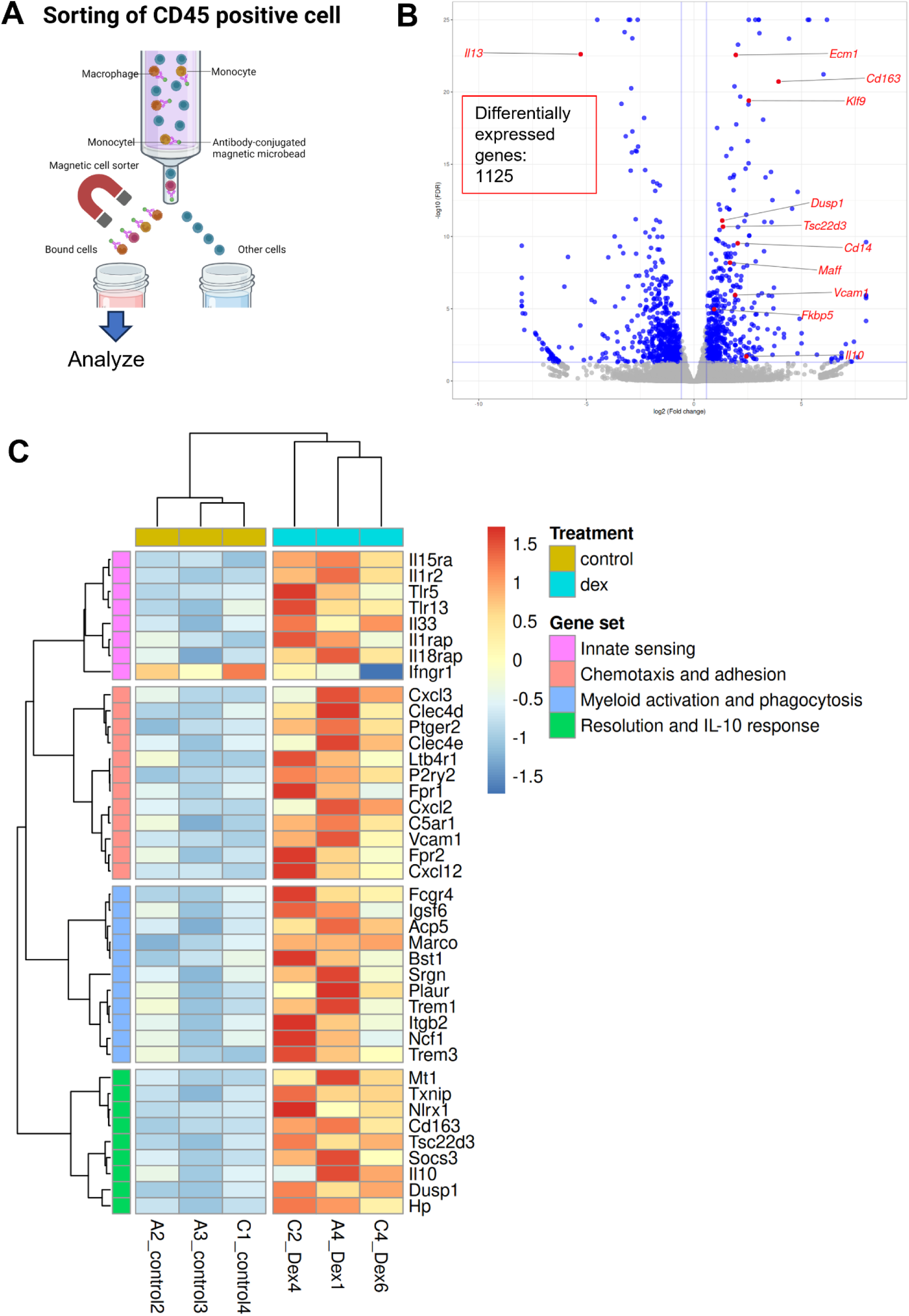
Dex treatment broadly affects immune cell gene expression and activates the IL-10 pathway in CD45⁺ cells. (A) Workflow for magnetic sorting (MACS) of E18.5 fetal testicular CD45⁺ cells. (B) Volcano plot of DEGs in MACS-purified immune cells. Major immune-related genes are highlighted in red. (C) Heatmap for pathway analysis showing IL10 axis/pathway activation. (D) Heatmap for module enrichment analysis of various pathways.

### Dex treatment induces the appearance of CD163^+^ macrophages in the fetal testis

We observed that CD163⁺ macrophage number increased after prenatal Dex treatment. While CD163⁺ cells were originally present only in the mesonephros in control fetuses, their numbers within the testis markedly increased following Dex exposure (Fig. 7A, 7C, 7D and 7E). CD163⁺ cells were GR⁺ and Ki67⁺ (Fig. 7A, 7C) and were observed in both testes and mesonephroi (Fig. 7B, 7E). No changes were observed in expression of the mitotic marker phospho-Histone H3 (pHH3) (Fig. 7D).

**Figure 7.**
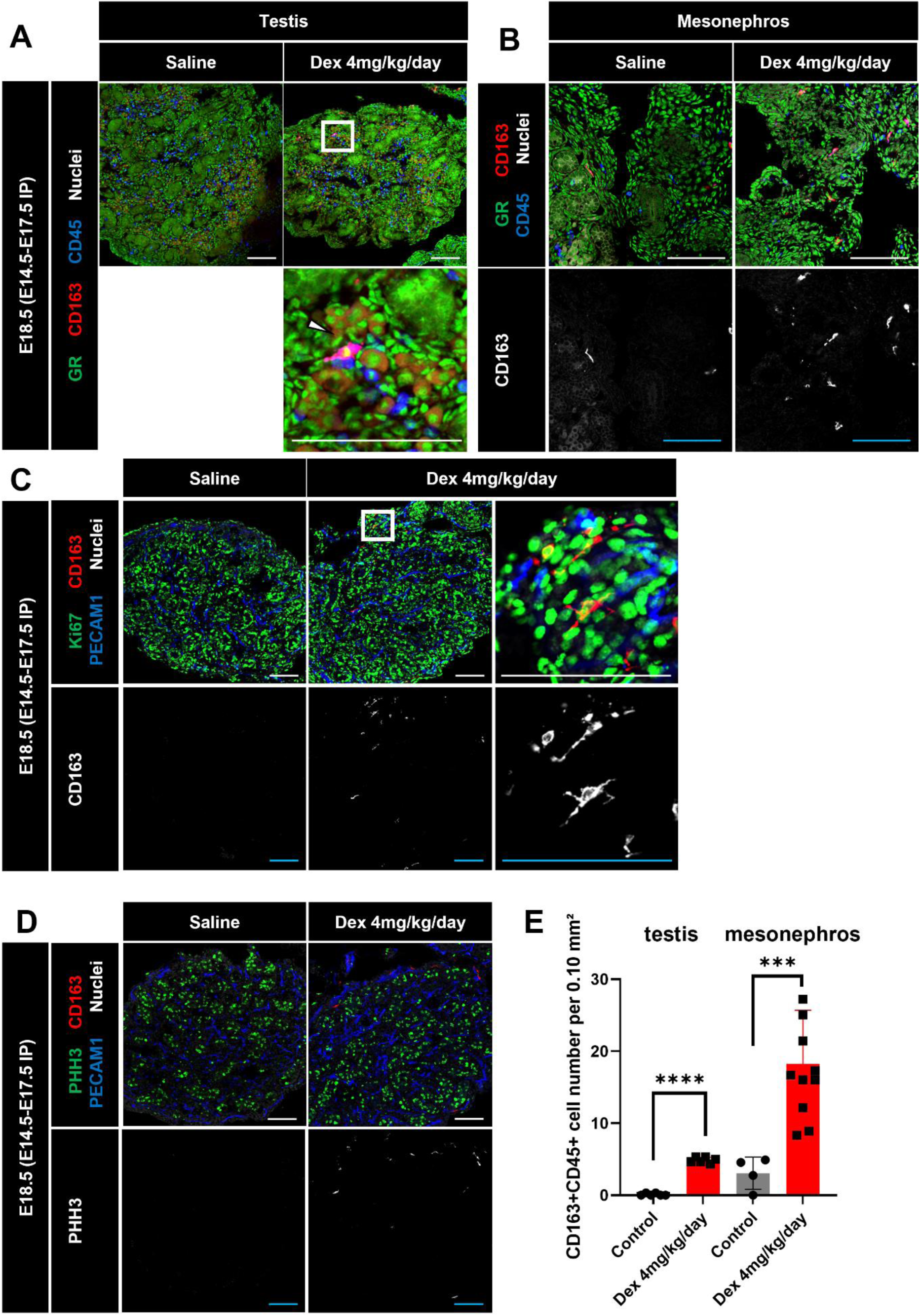
Prenatal Dex treatment increases CD163⁺ macrophage number in fetal testis and mesonephros. (A, B) Immunostaining for CD163 and CD45 in E18.5 testis and mesonephros, showing increased CD163 staining in Dex-treated versus saline control samples. (C) Ki67/CD163 testis staining. (D) pHH3/CD163 testis staining. (E) Quantification of CD163⁺CD45⁺ cells per unit area of testis and mesonephros. Scale bar: 100 μm. \**P*<0.05; \*\**P*<0.01; \*\*\**P*<0.001.

### IL10 induces hypervascularization and reduces Leydig cells

To assess the role of IL10 in fetal testis development and function, E12.5 and E13.5 testes were cultured ex vivo with IL-10 for 48 hours. In E12.5 testes, the number of CYP11A1-positive Leydig cells decreased in IL10-treated testes, while PECAM1⁺ endothelial cells increased (Fig. 8A, 8B, Fig. S12A). No significant changes were observed in E13.5 testes (Fig. 8A, Fig. S13B). pHH3 and Ki67 expression remained unchanged (Fig. S13A, S13B), and CD163⁺ cells were not detected following IL10 treatment (Fig. S13C). qRT-PCR confirmed significant upregulation of *cadherin 5* (*Cdh5*), an endothelial-cell-specific gene (Fig. 8C). Thus, these data suggest that IL10 increases endothelial cells and induces hypervascularization of the fetal testis (Fig. 9).

**Figure 8.**
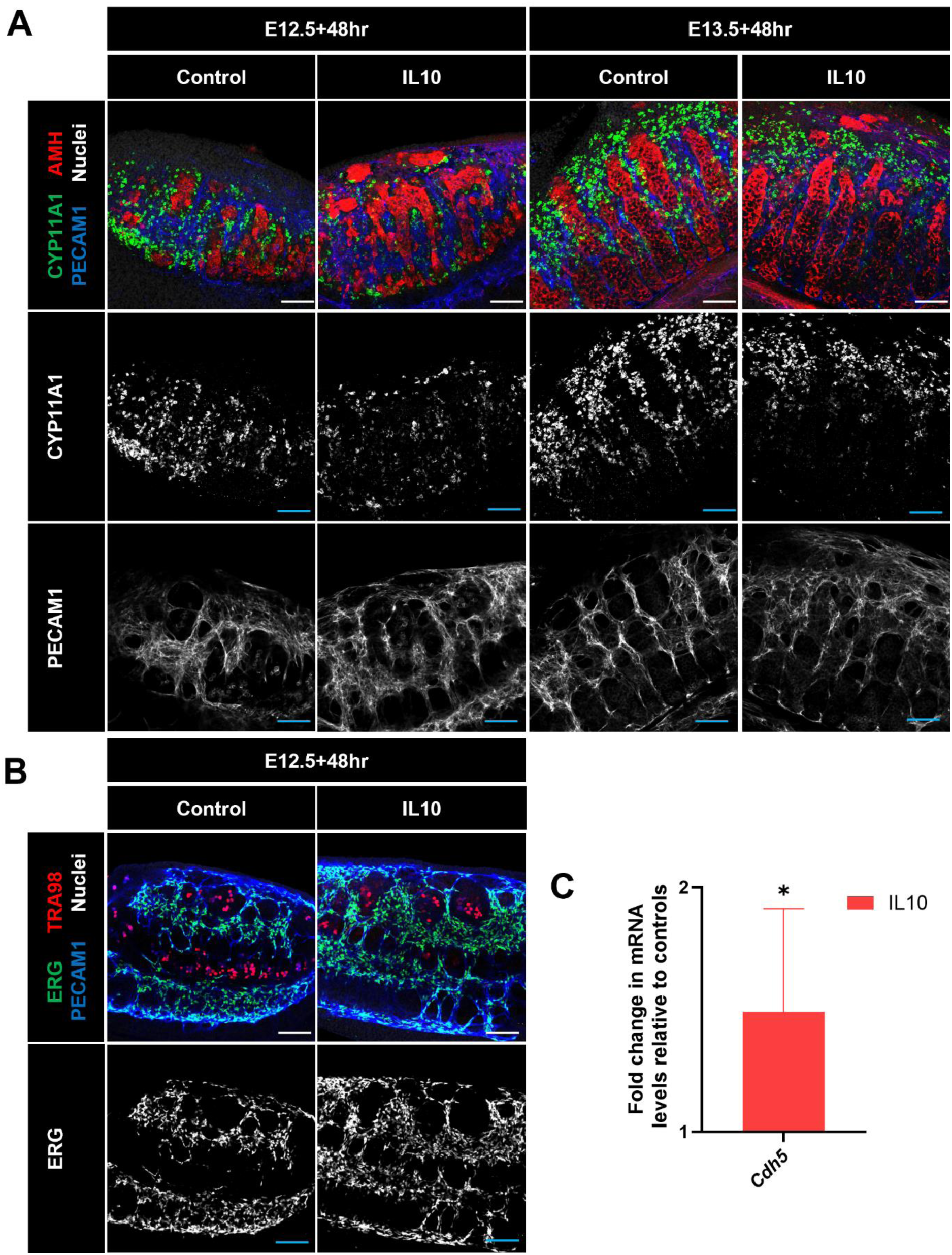
IL10 promotes endothelial expansion and reduces Leydig cells in ex vivo organ culture. (A) Immunostaining for CYP11A1 and PECAM1 in E12.5 and E13.5 testes cultured for 48 hrs with IL10-containing versus control media. (B) ERG staining showing endothelial cell nuclei. (C) qRT-PCR analysis of *Cdh5* (endothelial cell gene) expression. Scale bar: 100 μm. \**P*<0.05.

**Figure 9.**
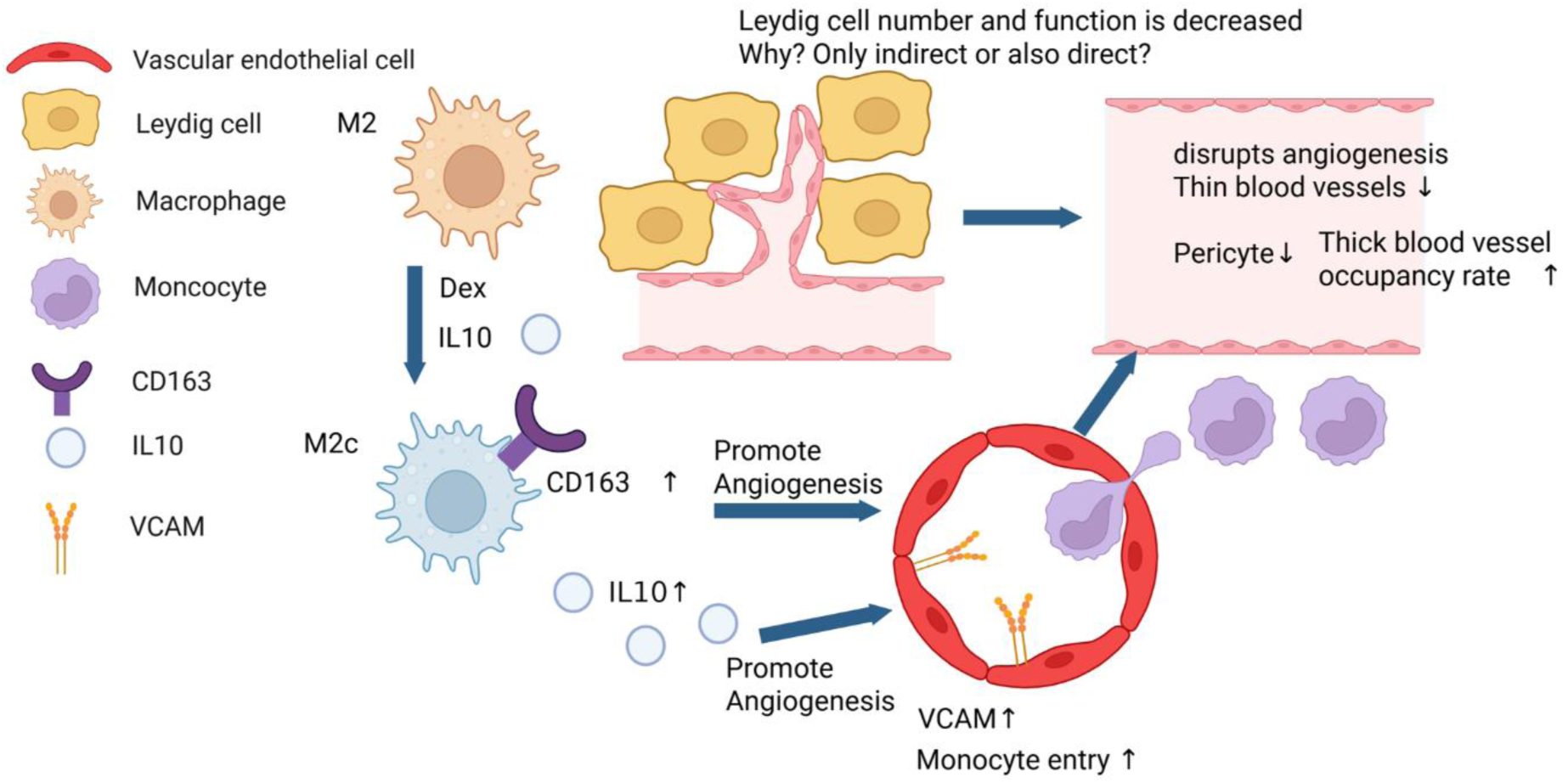
IL10-mediated vascular remodeling underlies testicular impacts of Dex treatment. Schematic model of Dex-induced IL10-mediated vascular remodeling.

## Discussion

Our research demonstrates that prenatal dexamethasone exposure suppresses androgen synthesis in the fetal testis by disrupting vascular-immune interactions, causing notable and specific alterations in fetal testis development. By combining immunofluorescence, 3D imaging, and transcriptome analyses, we observed that late-pregnancy glucocorticoid treatment activates immune programming and IL10 signaling, resulting in the induction and/or recruitment of CD163⁺ M2c macrophages. Furthermore, we demonstrated that such alterations in the immune environment cause vascular structural disruption and are linked to a reduction in Leydig cell number. Conversely, prenatal dexamethasone exposure did not exert detectable effects on ovarian gene expression, indicating an organ-specific and sex-dependent response to antenatal corticosteroid (ACS) therapy.

### Prenatal glucocorticoid exposure affects fetal immune and endocrine organs, including the testes

ACS administration is widely recognized as a fundamental intervention that improves preterm infant survival by promoting lung maturation (2, 3). However, the systemic effects of ACS on developing organs other than the lungs remain largely unclear. Indeed, it has been reported that prenatal betamethasone can remodel the developing immune system in the pancreas (4). Furthermore, not only ACS therapy but also maternal psychological stress, anxiety, and depression during pregnancy are consistently associated with increased fetal glucocorticoid exposure to both mother and fetus, mediated by decreased placental 11β-HSD2 activity and increased cortisol transfer to the fetus (27–29). Such increased fetal cortisol exposure has been reported to be associated with changes in infant temperament, cognitive development, and subsequent stress reactivity (30). Combined with evidence from prospective studies linking maternal psychosocial stress, anxiety, or depression during pregnancy to adverse offspring outcomes (e.g., altered neurodevelopment and increased risk of later psychopathology), psychosocial stress during pregnancy converges with prenatal corticosteroid exposure via common glucocorticoid pathways, potentially further sensitizing fetal immune and endocrine organs, including the testes (27, 30).

Regarding immune-mediated programming by prenatal glucocorticoids in the testes, studies primarily using rat models have suggested that prenatal glucocorticoid exposure may impair adult testicular structure and function (including reduced testicular weight and sperm count) (19), but the underlying mechanisms were not fully elucidated. More recently, Liu et al. showed that dexamethasone-activated GR in fetal Sertoli cells increases GR occupancy at the *Tbx2* promoter and recruits the coactivator p300, leading to increased H3K9ac at the TBX2 locus and upregulation of *Tbx2* (20). Increased TBX2 levels were proposed to suppress the expression of connexin 43 (CX43, also known as GJA1) expression, thereby compromising the function of the blood-testis barrier and ultimately contributing to reduced sperm quality. These epigenetic changes persist postnatally (20).

### Prenatal glucocorticoids suppress androgen synthesis in fetal testes via disruption of vascular-immune interactions

Our study demonstrates that prenatal glucocorticoid treatment can suppress androgen synthesis in the fetal testis via disruption of vascular-immune interactions, providing a mechanistic framework for interpreting these results. Our data shows that Dex exposure activates IL10 signaling within CD45⁺ immune cells, resulting in an increase of CD163⁺ M2c macrophages.

Previous studies have shown that the transmembrane scavenger receptor CD163 is expressed specifically in monocytes (low expression) and macrophages (high expression) (31). CD163⁺ M2/M2c macrophages promote angiogenesis and vascular permeability in inflammatory environments via the HIF1A/VEGFA pathway (23). Furthermore, in a Kazakh cohort of esophageal squamous cell carcinoma, CD163⁺ M2 macrophages were reported to contribute to tumor invasiveness and progression via matrix metallopeptidase 9 (MMP9) and angiogenesis (32). CD163⁺ macrophages are recognized as important immunoregulatory cells in many pathologies, including inflammation and tumors, and have also been shown to have potential as therapeutic targets (33). Furthermore, in a retinal ischemia model, *Il10*-deficient mice showed a marked reduction in pathological retinal angiogenesis (34). IL10 promotes retinal angiogenesis by altering macrophage angiogenic functions (34), consistent with the IL10-dependent changes observed in this study. Furthermore, it has been reported that IL10 overexpression induces increased vascular structure and epithelial re-epithelialization not only via macrophages but also through the mobilization of endothelial progenitor cells and increased VEGF expression (35).

Vascular cell adhesion molecule 1 (VCAM1) plays a crucial role in leukocyte adhesion and infiltration, tissue remodeling, and metastasis in pathologies such as rheumatoid arthritis, asthma, transplant rejection, and cancer (36). VCAM1 on endothelial cells not only functions as a scaffold for leukocyte migration but also induces endothelial cell signaling via reactive oxygen species (ROS) derived from NADPH oxidase (37). VCAM1-dependent monocyte/lymphocyte recruitment is also essential for lesion formation in classical vascular inflammation, such as atherosclerosis (38). Based on the above, Dex likely shifts fetal testicular macrophages toward an M2c-like (CD163-positive/IL10-producing) phenotype, thereby increasing VCAM1 expression on endothelial cells and promoting monocyte recruitment.

Previous studies have revealed that macrophages and blood vessels are not merely passive bystanders but actively orchestrate fetal testis morphogenesis. Yolk-sac-derived macrophages align along developing testicular blood vessels and are essential for establishing characteristic testicular vascular patterning and properly segmented testis cords. Depletion of these macrophages disrupts vascular patterning, resulting in malformed and fused testicular cords (39). Furthermore, disruption of angiogenesis and Notch signaling destabilizes this perivascular niche, promoting premature and unbalanced differentiation in multipotent cells and altering stromal architecture (40). Moreover, Li et al. showed that the skewed hematopoietic cell ratio caused by compound deficiency of *Mafb* and *Maf* leads to excessive production of myeloid immune cells, abnormal vascular patterning, and failure to form cords, demonstrating that not only the presence of myeloid subsets but also their identity and balance are important for establishing a normal vascular-stromal scaffold in the fetal testis (41). Furthermore, Gu et al. showed that yolk-sac-derived macrophages are responsible for the morphogenesis of blood vessels and testis cords in the fetal testis, while HSC-derived monocytes are recruited to the testes during a limited fetal time window and differentiate into interstitial macrophages and peritubular macrophages in adulthood, with interstitial macrophages specifically promoting steroid production in Leydig cells (24). Furthermore, excessive CD11b⁺ monocytes in the fetal testis have been shown to adversely affect testis cord morphogenesis and development, indicating that the ratio of monocytes to macrophages is more critical than the total macrophage number (41).

We found that Dex treatment skews fetal testicular macrophages toward an M2c-like phenotype and leads to an increased number of CD163⁺ macrophages. Given that CD163⁺ macrophages have been implicated in pro-angiogenic activity and that M2c macrophages secrete IL10, this shift is consistent with an IL10-driven pro-angiogenic/immunoregulatory program. In parallel, VCAM1 expression on endothelial cells increased, consistent with enhanced monocyte entry. Despite these signals, angiogenesis became dysregulated, characterized by a reduction in thin microvessels, decreased pericyte coverage, dilation of residual vessels, and a relative increase in thicker vessels. Consistent with our RNA-seq data showing reduced *Angpt2* expression, decreased *Angpt2* may alter angiopoietin-TEK (TIE2) signaling in a manner that favors vessel stabilization and limits vascular remodeling. As a result, fine new sprouts may fail to expand, and larger vessels may occupy a greater proportion of the vascular network. This altered vascular patterning was associated with reduced Leydig cell numbers and impaired steroidogenic function. Placed within the framework established by our previous work, these findings suggest that glucocorticoid exposure qualitatively reprograms macrophage-endothelial cell interactions and the behavior of the perivascular Nestin⁺ progenitor cell niche, rather than merely increasing macrophage or monocyte numbers. The timing of monocyte recruitment to the testes reported by Gu et al. (24) coincides with the period of Dex-induced monocyte increase observed in our study, suggesting that physiological monocyte recruitment may be amplified by ACS. Under normal conditions, yolk-sac-derived macrophages appear to promote the formation of a finely branched microvascular network that supports the balance between maintaining precursor cells and their differentiation into Leydig cells and smooth muscle cells. Under prenatal Dex administration, we propose that a set of imbalanced angiogenesis/remodeling signals is generated, characterized by chronically activated adhesive vascular surfaces, reduced capillaries, dilated large vessels, and loss of perivascular support, by skewing macrophages toward an M2c/IL10/VCAM1-associated state. Consequently, perivascular progenitor cells likely lose their stable niche, impairing the formation and maintenance of steroid-producing Leydig cell compartments. Indeed, ex vivo IL10 treatment induced marked alterations in vascular architecture. The Dex model we used in previous assays is based on E18.5 testes, so these E12.5 and E13.5 experiments are not directly comparable; however, these findings demonstrate that IL10 can influence gonadal development. At E12.5 and E13.5, yolk-sac-derived macrophages predominate in the testes, while fetal HSC-derived monocytes become the primary source of immune cells by E18.5. Therefore, it is possible that CD163-positive cells could not appear in IL10 culture experiments due to differences in the origin(s) of immune cells. Thus, while a reduction in Leydig cell numbers is thought to result from an indirect effect, the presence or absence of a direct effect on Leydig cells requires further verification. Leydig cells express GR and are thus susceptible to direct glucocorticoid action. Furthermore, in vitro studies suggest that Dex inhibits Leydig cell differentiation, and this effect is reversible with NR3C1 (GR) antagonists (42).

### Prenatal GC therapy induces organ-specific and sex-dependent responses

This study revealed that prenatal Dex exposure did not exert any detectable effects on ovarian gene expression, indicating the existence of organ-specific and sex-dependent responses to ACS therapy. Previous studies have reported that both testicular and ovarian germ cells exhibit dynamic temporal regulation of GR expression during fetal and adult development, with patterns clearly differing by sex (43). Our study demonstrated that developmental GR expression patterns differ not only in germ cells but also in other cell types. Furthermore, the response following Dex administration also differed significantly between testes and ovaries, with no marked effect observed in ovaries. This sex-specific phenotype may reflect differences in the fetal gonadal chromatin landscape and GR signaling competence between testes and ovaries, which can influence GR genomic binding and downstream transcriptional responses in a tissue-dependent manner. Future studies that directly profile GR binding and chromatin accessibility in fetal gonads will be required to define the mechanisms underlying this sex-biased response.

### GR expression in the human testis

Although this study used mice, it has been reported that in the adult human testis, GR is detected in peritubular cells, a subset of Leydig cells, Sertoli cells (weakly), and spermatogonia, but not in spermatocytes (44). The GR expression pattern in fetal testis samples differs, showing heterogeneous expression particularly in Sertoli cells, absent expression in spermatogonia, and weak expression in newly formed peritubular cells. Furthermore, GR expression is detected in a subset of pro-spermatogonia (45). Epidemiological studies in humans are anticipated to provide future insights into the extent to which these molecular mechanisms translate into clinical outcomes.

### Limitations

Several limitations should also be acknowledged. First, this study was conducted in a mouse model, requiring caution when extrapolating findings to human physiology. Second, relatively high doses of Dex were used to achieve pharmacokinetic equivalence with human ACS regimens, and the dose-response relationship remains incompletely elucidated. Third, this analysis was limited to late pregnancy and did not include long-term assessment of adult reproductive capacity or endocrine function. Finally, while bulk RNA-seq provided valuable insights into transcriptional changes, single-cell analysis techniques would help elucidate cell-type-specific responses and detailed signaling dynamics. Future work should validate these findings in human settings through more refined analysis of fetal tissues and longitudinal studies of individuals exposed to ACS in utero.

## Methods

### Animals

C57BL/6J mice (JAX stock #000664) were used for all experiments. All animals were housed in accordance with National Institutes of Health guidelines. Experimental protocols were approved by the Institutional Animal Care and Use Committee (IACUC) of Cincinnati Children’s Hospital Medical Center (protocol numbers: IACUC2021-0016 and IACUC2024-0039).

### Gonad culture

E12.5 and E13.5 testes were cultured ex vivo using an agar-based culture method as previously described (46). IL10 (Recombinant Murine IL-10; PeproTech #210-10-10μg) was diluted in 1% bovine serum albumin (BSA) to a final concentration of 100 ng/mL or 500 ng/mL. Testes treated with an equivalent volume of 1% BSA served as negative controls. After 48 hours of culture, samples were collected for whole-mount immunofluorescence or RNA extraction. A minimum of three independent experiments were conducted, with each experiment using multiple testes (*n*=3-5).

### Immunofluorescence

Samples were fixed overnight at 4°C in phosphate-buffered saline (PBS) containing 4% paraformaldehyde (PFA) and 0.1% Triton X-100. Whole-mount immunofluorescence was performed on testes at embryonic stages E11.5, E12.5, and E13.5. All other samples (E14.5 and older) were analyzed via cryosection immunofluorescence.

Following multiple washes in PBTx (PBS with 0.1% Triton X-100), testes were blocked for 1 hour at room temperature (RT) using a solution of 10% fetal bovine serum (FBS) and 3% bovine serum albumin (BSA). Primary antibodies were diluted in blocking solution and incubated overnight at 4°C on a shaking platform. After additional PBTx washes, samples were incubated with fluorescently labeled secondary antibodies for 1 hour at room temperature with gentle agitation. For whole-mount samples, the incubation time for secondary antibodies was extended to 3 hours. Final washes in PBTx were followed by mounting with Fluoromount-G (Southern Biotech).

Details of primary antibodies and dilutions used are provided in Supplementary Table S1. Secondary antibodies conjugated to Alexa Fluor 488, Alexa Fluor 555, and Alexa Fluor 647 (Molecular Probes/Life Technologies/Thermo Fisher) were used at a dilution of 1:500. Nuclear staining was performed using Hoechst 33342 at 2 μg/mL (#H1399, Thermo Fisher) and is referred to as “Nuclei” in all image labels. Microscopy was performed using a Nikon A1 inverted LUNV microscope (Nikon, Tokyo, Japan), operated with NIS-Elements software (PerkinElmer, Nikon, Tokyo, Japan).

### 3D whole-mount immunofluorescence

Samples were fixed in 4% PFA prepared in PBTx overnight at 4°C on a rocker. Following fixation, tissues were rinsed twice in PBTx at RT and subsequently washed three times for 10 min each in PBTx at RT on a rocker. Lipid, heme, and lipofuscin were extracted by incubating tissues in a solution containing 25% Quadrol and 5% CHAPS for 1 day at RT on a rocker. Samples were then rinsed twice in PBTx at RT and washed three times for 10 min each in PBTx at RT on a rocker. Blocking was performed for 1 day at RT on a rocker using blocking buffer composed of 0.5% Triton X-100, 0.5% CHAPS, 10% fetal bovine serum (FBS), and 3% BSA. Tissues were incubated in primary antibody solution prepared in the same blocking buffer for 3 days at 37°C on a rocker. After incubation, samples were rinsed twice in PBTx at RT and washed three times for 10 min each and overnight in PBTx at RT on a rocker. Tissues were incubated in secondary antibody for 2 days at 37°C. After incubation with secondary antibodies, tissues were washed for 1 day. For mounting, tissues were transferred into a MatTek coverglass-bottom dish. Wash buffer was removed and replaced with a small amount of EZ View mounting medium (7M urea, 80% Nycodenz in 0.02M phosphate buffer). A 22 mm cover glass was placed on top to weigh down the tissue and prevent evaporation. Imaging was performed immediately after mounting using a Yokogawa_W1 spinning disk confocal system on a Nikon Ti2 microscope. Analyses were performed using Bitplane Imaris 10.2. Vessel volumes were measured using AI surface segmentation.

### RNA extraction and quantitative real-time PCR (qRT-PCR)

Gonads were carefully separated from the mesonephros, and total RNA was extracted using TRIzol reagent (Invitrogen/Thermo Fisher) according to the manufacturer’s protocol. Complementary DNA (cDNA) was synthesized using the iScript cDNA synthesis kit (Bio-Rad), with 500 ng of total RNA per reaction.

qRT-PCR was performed using the Step OnePlus Real-Time PCR System (Applied Biosystems/Thermo Fisher) with Fast SYBR Green Master Mix (Applied Biosystems/Thermo Fisher). The thermal cycling conditions were: 95°C for 20 seconds, followed by 40 cycles of 95°C for 3 seconds and 60°C for 30 seconds. *Gapdh* was used as the reference gene for normalization. Relative gene expression was calculated using the ΔΔCt method, and data are presented as mean ± standard deviation (SD) from experiments including at least three gonads per biological replicate. Primer sequences are listed in Supplementary Table S2.

### RNA extraction and bulk RNA sequencing (RNA-seq)

Gonads were separated from the mesonephros, and total RNA was extracted using TRIzol reagent (Invitrogen/Thermo Fisher). For each sample, four testes and eight ovaries from the same dam were pooled into a single tube for RNA extraction. All samples were collected from fetuses of the same dam.

### Bulk RNA-seq analysis

All processing of RNA-seq used the Ensembl (47) *Mus musculus* reference genome (GRCm39/mm39) and the corresponding gene annotation. Raw paired-end reads underwent quality control and adapter trimming with fastp v0.23.2 (48). Reads passing quality control were aligned to the reference genome with HISAT2 v2.2.1 (49). Aligned reads were converted to Binary Alignment/Map (BAM) format with SAMtools v1.21.0 (50), sorted with BamTools v2.5.1(51), and quantified at the gene level using StringTie v2.2.1 (52).

### Differential Expression Analysis

Differentially expressed genes (DEGs) were detected using DESeq2 v1.48.2 (53) with default parameters, after removing genes without 10 raw read counts across all samples. Genes were considered significant at absolute fold change > 1.5 and false discovery rate (FDR) < 0.05 (54). Principal component analysis (PCA) was performed on the 500 genes with the largest absolute fold changes. For heatmap visualization, normalized counts by DESeq2 passed Z-score standardization and were plotted with pheatmap v1.0.13 (Kolde, 2025; R package, https://github.com/raivokolde/pheatmap). Hierarchical clustering used Ward’s minimum-variance method (55), applied separately to upregulated and downregulated gene sets.

### Gene set enrichment analysis (GSEA)

GSEA was performed using MSigDB (56) and fgsea v1.34.2 (57). Enrichment scores are estimated for hallmark gene sets, immunologic signature gene sets, and canonical pathways.

### Statistics

Details of statistical analyses, including sample size (*n*), definitions of *n*, measures of central tendency and variability (mean ± SD), and criteria for statistical significance are provided in the figure legends. For qRT-PCR data, statistical analyses were performed using Prism software version 10.0 (GraphPad), applying an unpaired, two-tailed Student’s *t*-test to ΔCt values. Each qRT-PCR experiment included a minimum of three biological replicates.

For immunofluorescence experiments, at least three sections from a minimum of three different animals (constituting three independent biological replicates) were analyzed per developmental stage or experimental condition. For cell counting and morphometric analyses, sample sizes for each group are explicitly stated. Graphical data are presented as mean ± SD, and statistical comparisons were made using an unpaired, two-tailed Student’s *t*-test.

## Acknowledgments

This work was supported by the National Institutes of Health (NIH) (grant R01HD094698 to T.D.), Uehara Memorial Foundation Fellowship (to S.M.), Japan Society for the Promotion of Science Overseas Research Fellowship (to S.M.), JSPS KAKENHI Grant Number 23K15775 (to S.M.), and Cincinnati Children’s Hospital Medical Center.

## Author Contributions

S.M and T.D. designed research; S.M., L.H., K.M., S.-Y.L., M.G., X.G., M.K., V.R., T.S., and T.D. performed research; S.M., S.-Y.L., K.M and T.D. analyzed data; S.M., L.H.K.M., S.-Y.L., , M.G., X.G., M.K., V.R., T.S., and T.D. edited the paper; and S.M., K.M., and T.D. wrote the paper.

