## Supplementary Data for "Prenatal corticosteroid exposure disrupts vascular-immune interactions and impairs steroidogenesis in the fetal testis"

**This file contains:**

Supplementary Figures 1-14
Supplementary Tables S1-S2
Supplementary References

### Supplementary Figures and Figure Legends

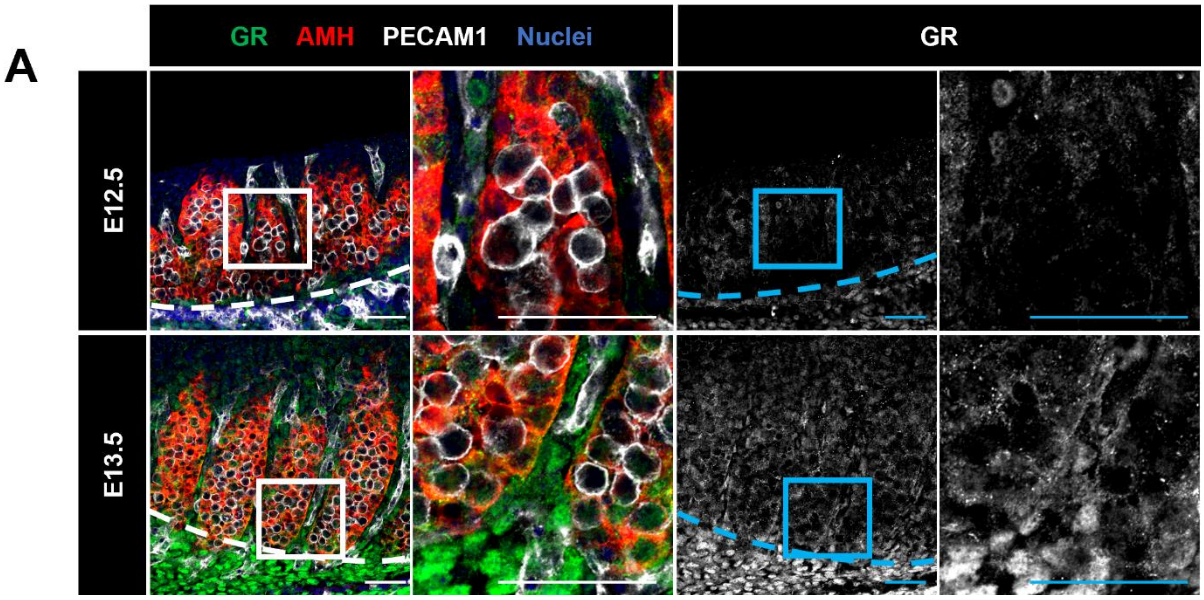


**Supplementary Figure 1. GR expression is limited in early fetal testes.**

(A) Immunofluorescence images of E12.5 and E13.5 XY gonads showing GR localization. Scale bar: 100 µm. Dashed lines: gonad–mesonephros border.

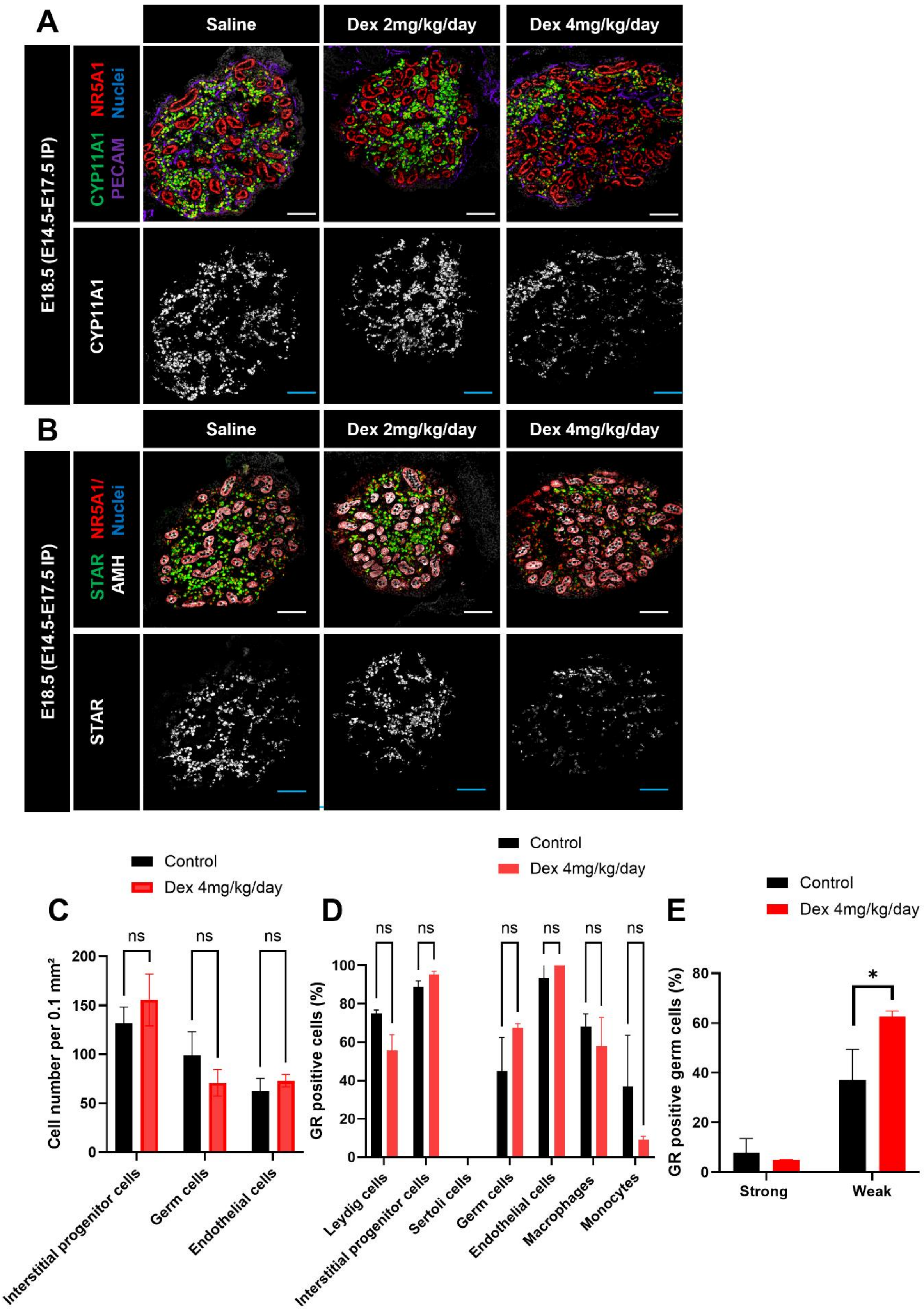
A
Saline
Dex 2mg/kg/day
Dex 4mg/kg/day
E18.5 (E14.5-E17.5 IP)
CYP11A1 NR5A1 PECAM Nuclei
CYP11A1
B
Saline
Dex 2mg/kg/day
Dex 4mg/kg/day
E18.5 (E14.5-E17.5 IP)
STAR NR5A1/ AMH Nuclei
STAR
C
Control
Dex 4mg/kg/day
Cell number per 0.1 mm²
ns
Interstitial progenitor cells
Germ cells
Endothelial cells
D
Control
Dex 4mg/kg/day
GR positive cells (%)
ns
Leydig cells
Interstitial progenitor cells
Sertoli cells
Germ cells
Endothelial cells
Macrophages
Monocytes
E
Control
Dex 4mg/kg/day
GR positive germ cells (%)
*
Strong
Weak

**Supplementary Figure 2. Prenatal Dex treatment suppresses androgen production.**

(A,B) Immunofluorescence images of E18.5 testes from saline control, Dex 2 mg/kg/day, and Dex 4 mg/kg/day groups. (C) Quantification of number of interstitial progenitors, germ cells, and endothelial cells per unit area (0.1 $mm^2$) of gonad in E18.5 saline-treated and Dex-treated testes. (D) Quantification of percent GR-positive Leydig cells, interstitial progenitors, Sertoli cells, germ cells, endothelial cells, macrophages, and monocytes in E18.5 saline control and Dex-treated testes. (E) Quantification of percent germ cells with strong versus weak GR expression in E18.5 control and Dex-treated testes. Scale bar: 100 µm. *$P$<0.05; ns: not significant.

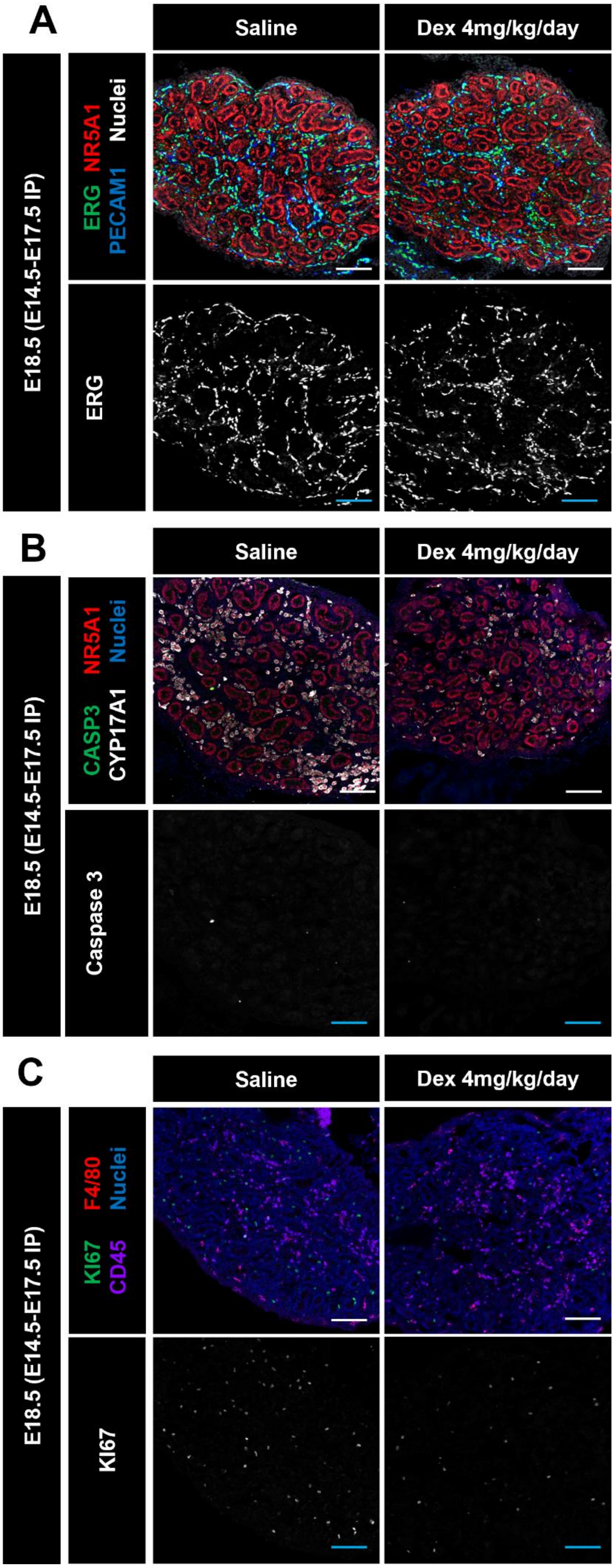
A
Saline
Dex 4mg/kg/day
E18.5 (E14.5-E17.5 IP)
ERG NR5A1 PECAM1 Nuclei
ERG
B
Saline
Dex 4mg/kg/day
E18.5 (E14.5-E17.5 IP)
CASP3 NR5A1 CYP17A1 Nuclei
Caspase 3
C
Saline
Dex 4mg/kg/day
E18.5 (E14.5-E17.5 IP)
KI67 F4/80 CD45 Nuclei
KI67

37

**Supplementary Figure 3. Cell proliferation and apoptosis after Dex treatment.**

(A) Immunostaining for CASP3/NR5A1/CYP17A1 in E18.5 saline control or Dex-treated fetal testes. (B) CASP3 and Ki67 staining. (C) Ki67/F4/80/CD45 staining to assess immune cell proliferation. Scale bar: 100 μm.

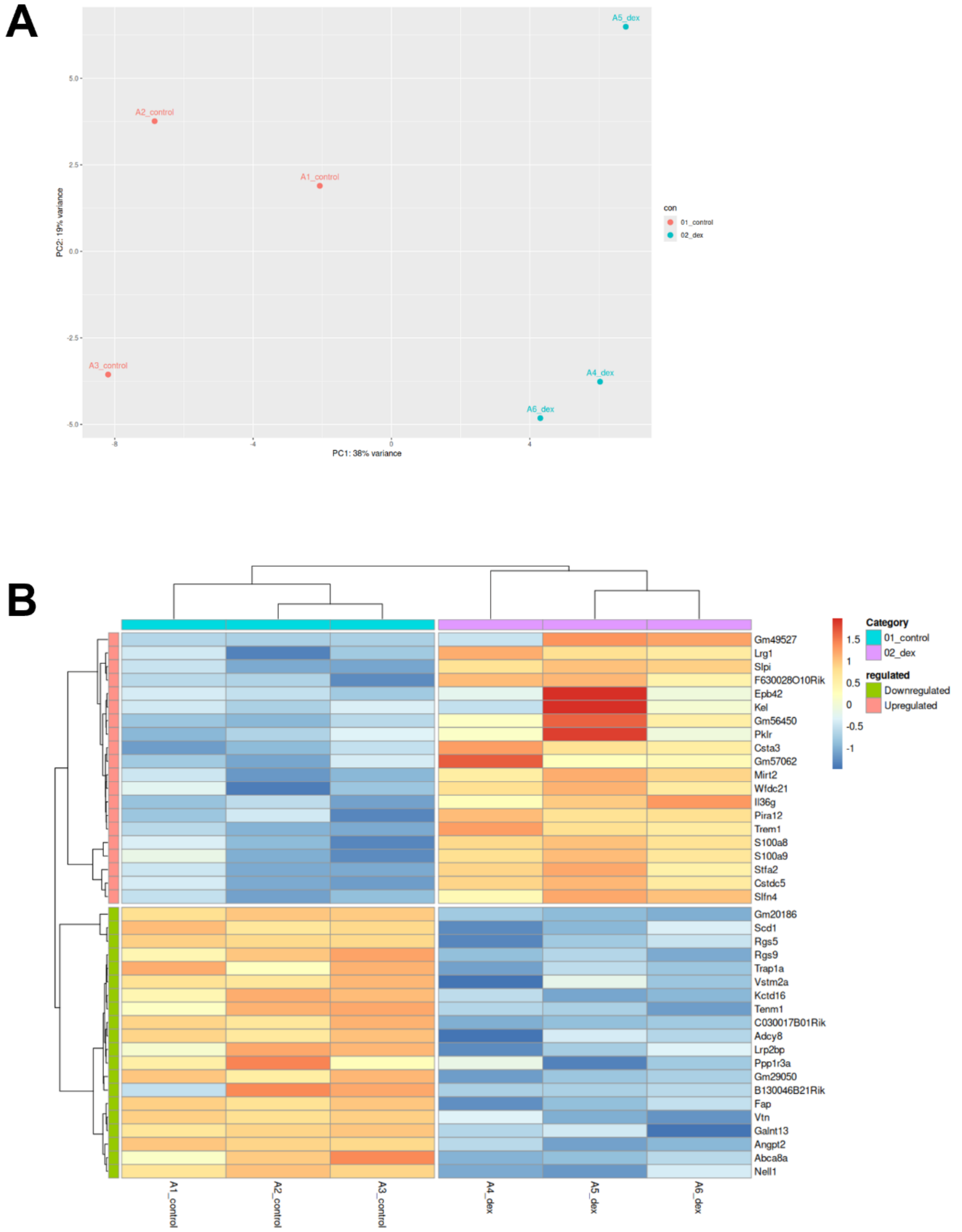


**Supplementary Figure 4. PCA and DEG distribution in testes after Dex treatment.**

(A) PCA plot of RNA-seq data showing separation between saline and Dex-treated samples. (B) Heatmaps of the top 20 differentially expressed genes.

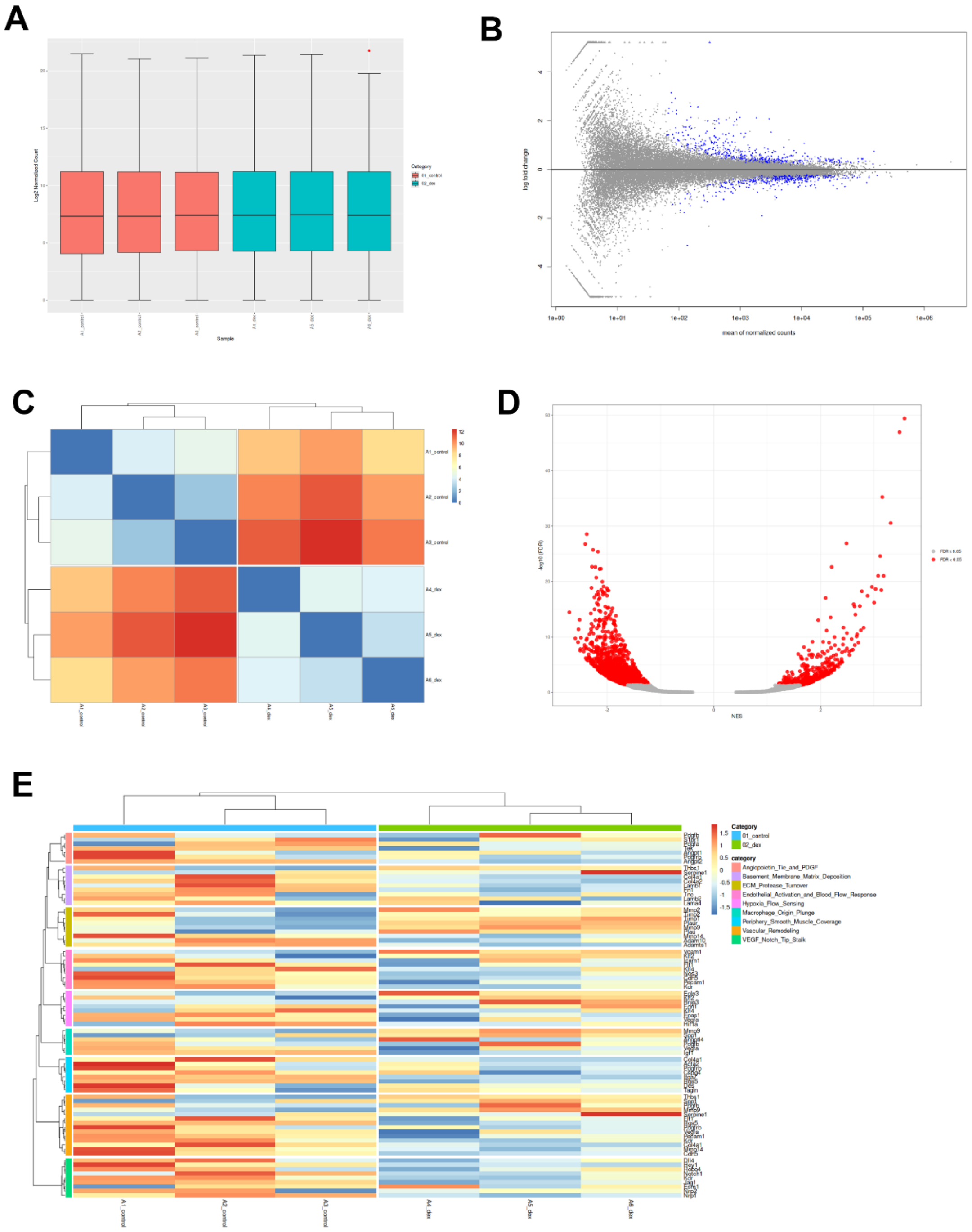


**Supplementary Figure 5. Transcriptomic analysis after Dex treatment.**

(A) Distribution of log2-normalized counts across samples (n = 3 control, n = 3 Dex). (B) MA plot of differential expression (Dex vs control). (C) Sample similarity heatmap with hierarchical clustering. (D) Pathway/gene-set enrichment volcano plot (NES vs –log10 FDR). (E) Heatmap of scaled expression of genes from enriched vascular pathways (endothelial activation, ECM turnover, angiogenesis, and vascular remodeling-related programs).

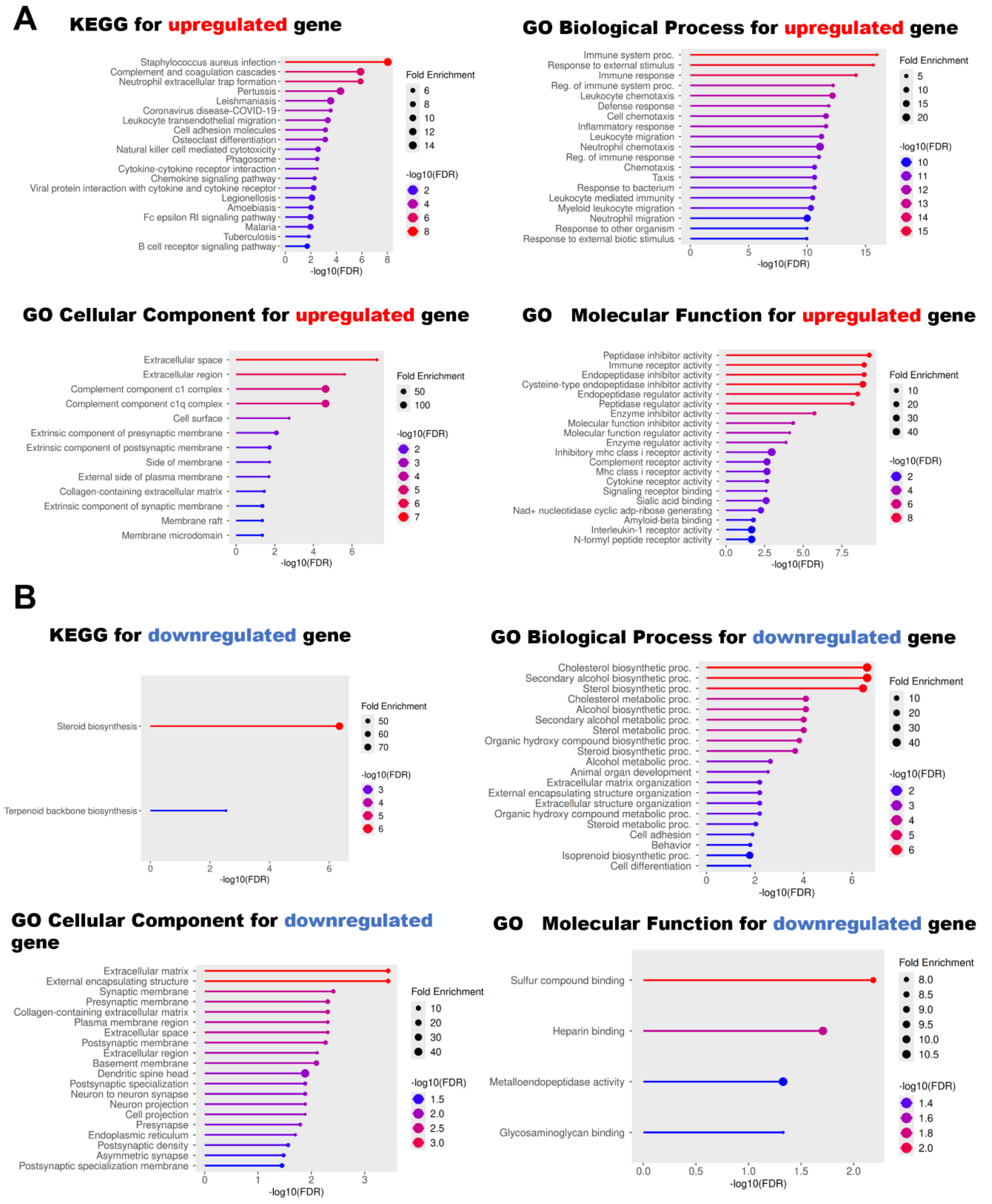


**Supplementary Figure 6. Functional enrichment of immune and steroidogenic pathways.** (A) KEGG and GO analyses for upregulated genes showing enrichment in immune response, chemotaxis, and inflammatory pathways. (B) KEGG and GO analyses for downregulated genes showing suppression of cholesterol and steroid biosynthesis.

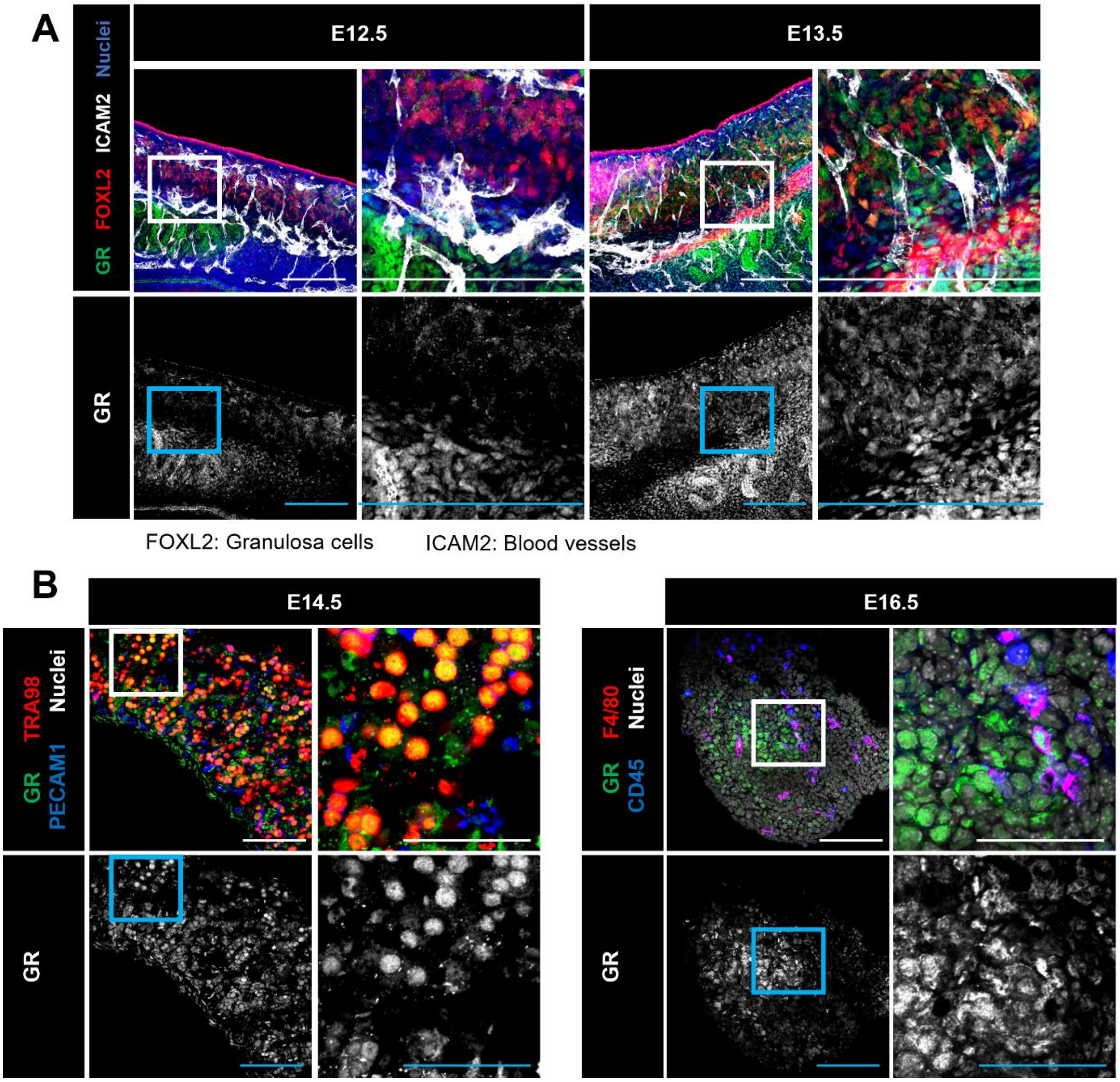


**Supplementary Figure 7. GR expression in fetal ovaries.**

(A) Immunostaining for GR/FOXL2/ICAM2 in E12.5 and E13.5 ovaries. FOXL2 marks granulosa cells; ICAM2 marks blood vessels. (B) GR/F4/80/CD45 staining in E16.5 ovaries. Scale bar: 100 µm.

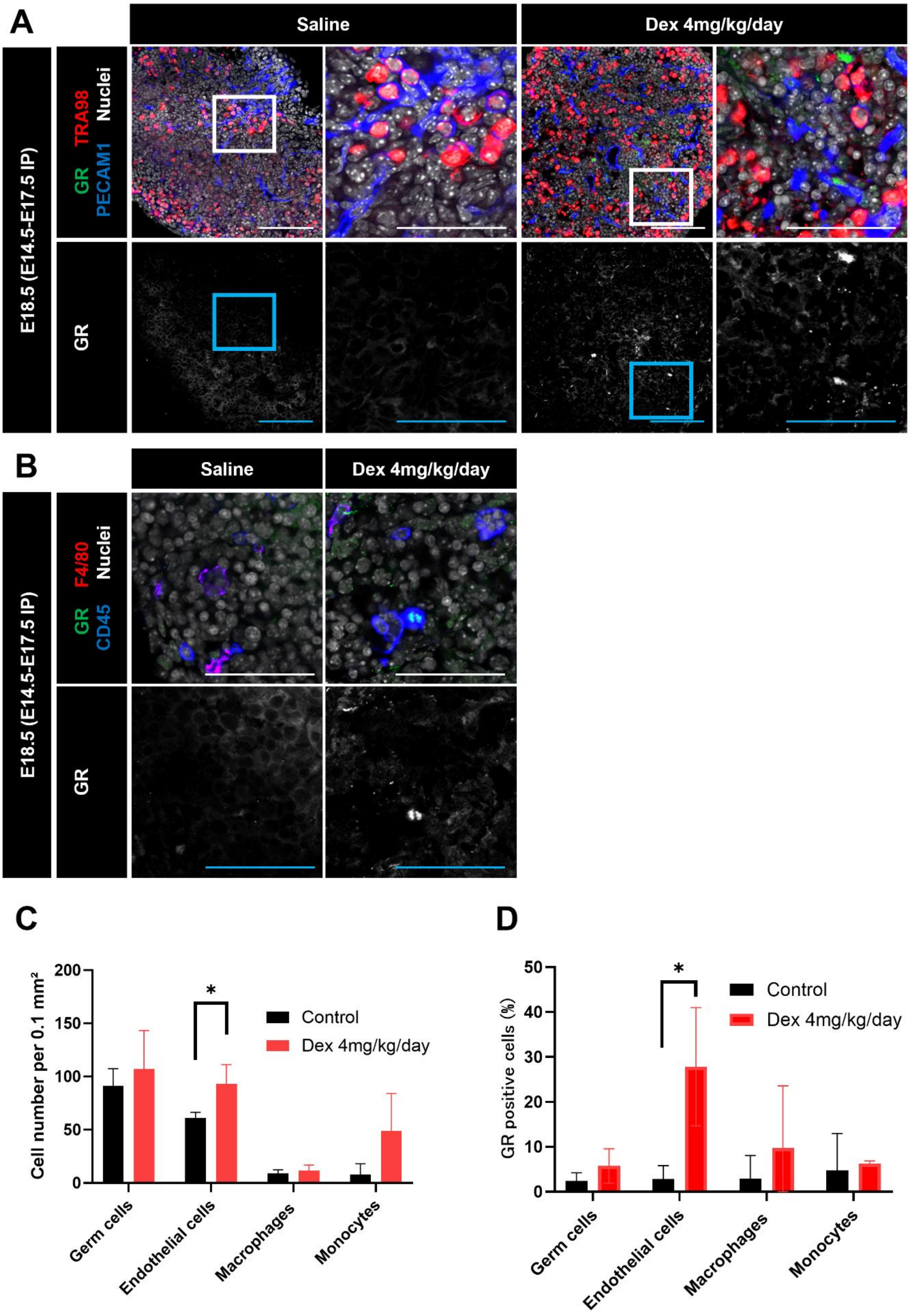
A
Saline
Dex 4mg/kg/day
E18.5 (E14.5-E17.5 IP)
GR TRA98 Nuclei
PECAM1
GR
B
Saline
Dex 4mg/kg/day
E18.5 (E14.5-E17.5 IP)
GR F4/80 Nuclei
CD45
GR
C
Cell number per 0.1 mm²
Control
Dex 4mg/kg/day
Germ cells
Endothelial cells
Macrophages
Monocytes
D
GR positive cells (%)
Control
Dex 4mg/kg/day
Germ cells
Endothelial cells
Macrophages
Monocytes

62

**Supplementary Figure 8. GR distribution after Dex treatment in ovaries.**

(A) Immunostaining for GR/TRA98/PECAM1 in E18.5 ovaries after saline or Dex treatment. (B) GR/F4/80/CD45 staining. (C) Quantification of GR-positive germ cells, endothelial cells, macrophages, and monocytes per unit area (0.1 $mm^2$) of ovary. (D) Quantification of percent GR-positive cell types E18.5 fetal ovaries after saline or Dex treatment. Scale bar: 100 µm. *$P<0.05$.

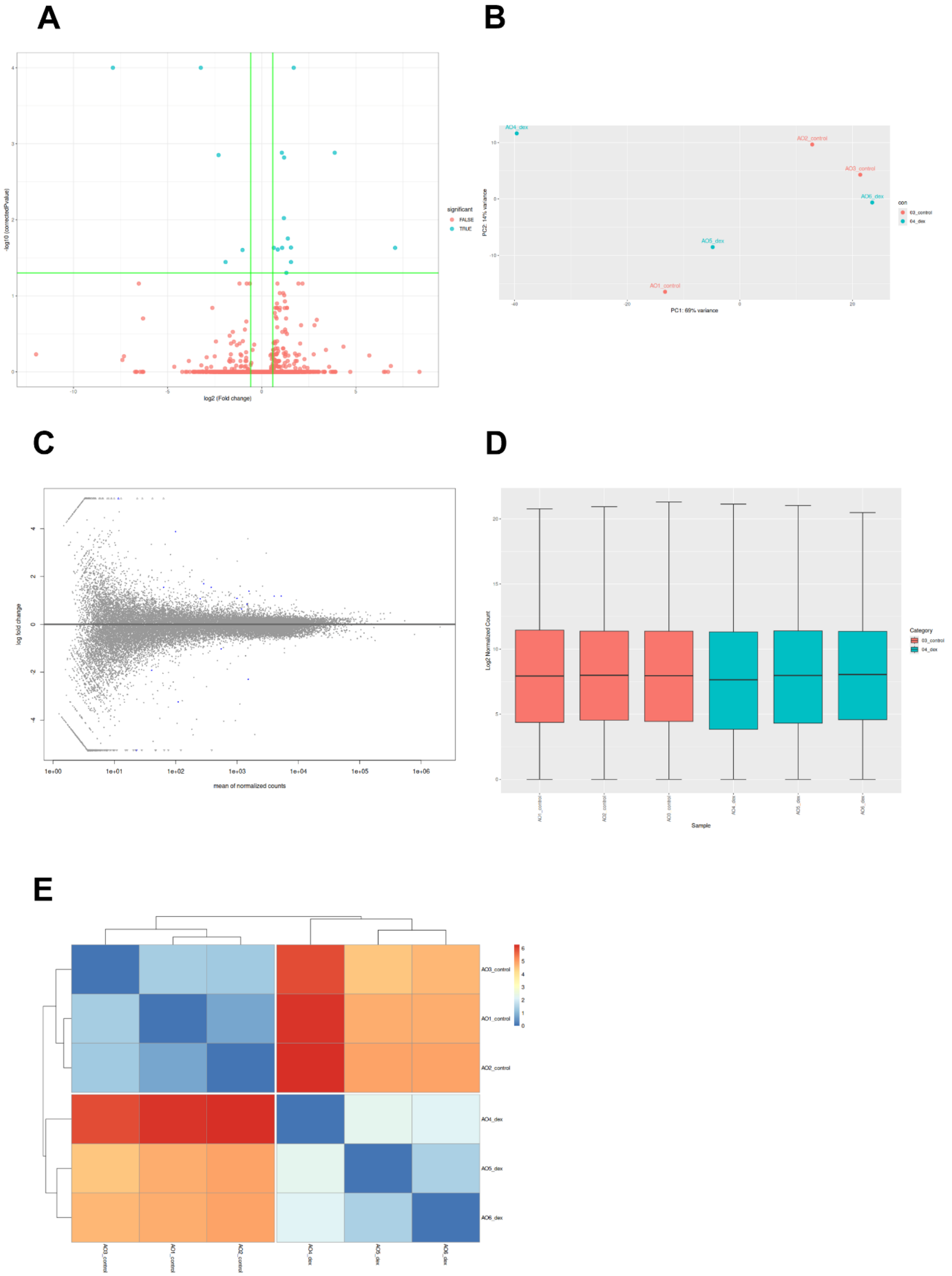
A
B
C
D
E
-log10 (correctedPvalue)
log2 (Fold change)
significant
FALSE
TRUE
PC2: 14% variance
PC1: 69% variance
AO4_dex
AO2_control
AO3_control
AO6_dex
AO5_dex
AO1_control
con
log fold change
mean of normalized counts
1e+00
1e+01
1e+02
1e+03
1e+04
1e+05
1e+06
Log2 Normalized Count
Sample
Category
03_control
04_dex
AO3_control
AO1_control
AO2_control
AO4_dex
AO5_dex
AO6_dex

**Supplementary Figure 9. Ovarian transcriptome analysis: PCA and differential gene expression.**

(A) Volcano plot of differential expression (Dex vs control; log2FC vs –log10 adjusted P value), showing few genes passing the significance/effect-size thresholds (green lines). (B) PCA of ovarian RNA-seq samples colored by treatment group. (C) MA plot summarizing log2 fold change versus mean normalized counts. (D) Box plots of log2-normalized count distributions across samples. (E) Sample-to-sample similarity heatmap with hierarchical clustering.

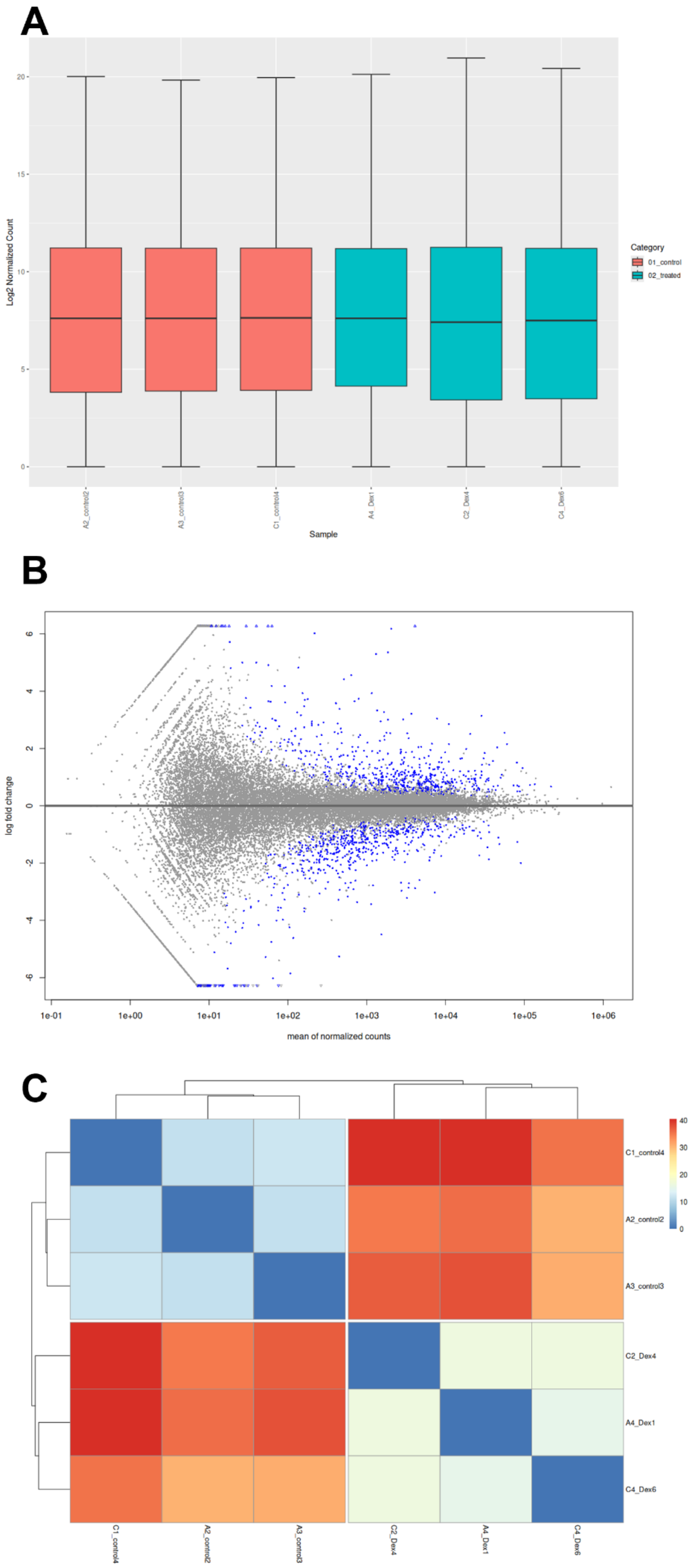
A
Log2 Normalized Count
Sample
Category
01_control
02_treated
A2_control2
A3_control3
C1_control4
A4_Dex1
C2_Dex4
C4_Dex6
B
log fold change
mean of normalized counts
1e-01
1e+00
1e+01
1e+02
1e+03
1e+04
1e+05
1e+06
C
C1_control4
A2_control2
A3_control3
C2_Dex4
A4_Dex1
C4_Dex6

**Supplementary Figure 10. Expression distribution and fold-change analysis in purified testicular CD45$^+$ cells.**

(A) Box plots showing the distribution of log2-normalized counts across RNA-seq samples from purified CD45$^+$ cells (3 control and 3 Dex-treated replicates), indicating comparable global expression distributions between groups. (B) MA plot of differential expression (Dex vs control), showing log2 fold change versus the mean of normalized counts; genes meeting the significance criteria are highlighted (blue). (C) Sample-to-sample similarity (distance) heatmap with hierarchical clustering based on global transcriptomic profiles, demonstrating separation of control and Dex-treated CD45$^+$ cell samples.

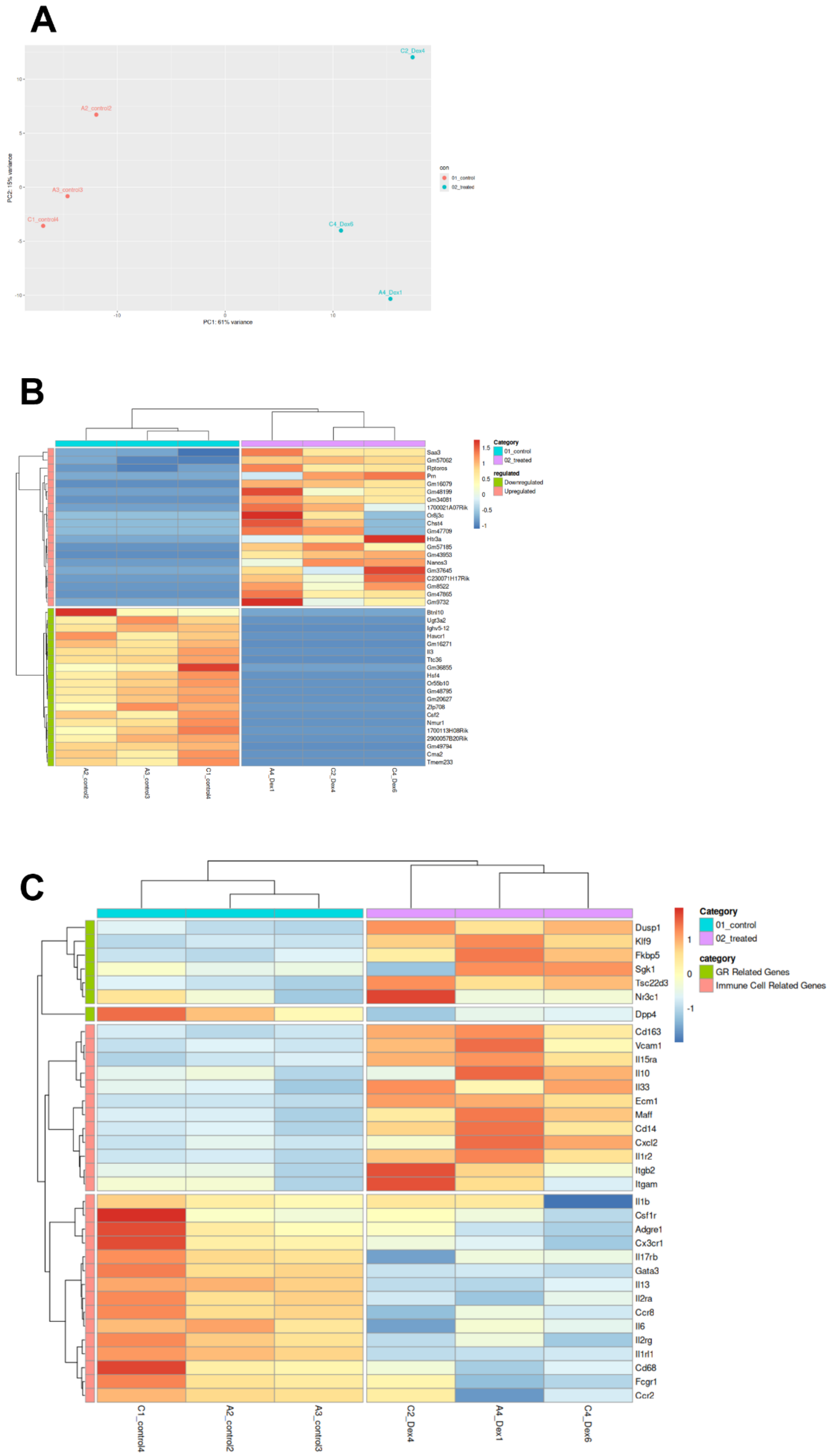
A
C2_Dex4
A2_control2
A3_control3
C1_control4
C4_Dex6
A4_Dex1
PC1: 61% variance
PC2: 15% variance
10
5
0
-5
-10
con
01_control
02_treated
B
Category
01_control
02_treated
regulated
Downregulated
Upregulated
1.5
1
0.5
0
-0.5
-1
Saa3
Gm57062
Rptoros
Prn
Gm16079
Gm48199
Gm34081
1700021A07Rik
Or8j3c
Chst4
Gm47709
Htr3a
Gm57185
Gm43953
Nanos3
Gm37645
C230071H17Rik
Gm8522
Gm47865
Gm9732
Btnl10
Ugt3a2
Ighv5-12
Havcr1
Gm16271
Il3
Ttc36
Gm36855
Hsf4
Or55b10
Gm48795
Gm20627
Zfp708
Csf2
Nmur1
1700113H08Rik
2900057B20Rik
Gm49794
Cma2
Tmem233
A2_control2
A3_control3
C1_control4
A4_Dex1
C2_Dex4
C4_Dex6
C
Category
01_control
02_treated
category
GR Related Genes
Immune Cell Related Genes
1
0
-1
Dusp1
Klf9
Fkbp5
Sgk1
Tsc22d3
Nr3c1
Dpp4
Cd163
Vcam1
Il15ra
Il10
Il33
Ecm1
Maff
Cd14
Cxcl2
Il1r2
Itgb2
Itgam
Il1b
Csf1r
Adgre1
Cx3cr1
Il17rb
Gata3
Il13
Il2ra
Ccr8
Il6
Il2rg
Il1rl1
Cd68
Fcgr1
Ccr2
C1_control4
A2_control2
A3_control3
C2_Dex4
A4_Dex1
C4_Dex6

**Supplementary Figure 11. GR-responsive and immune-related gene expression in purified testicular CD45$^{+}$ cells.**

(A) PCA Plot for top 500 DEGs. (B) Heatmap (for top 20 upregulated / downregulated DEGs). (C) Heatmap of GR-responsive genes (*Tsc22d3*, *Fkbp5*) and immune cell markers (*Cd163*, *Vcam1*, *Itgam*) in testicular CD45$^{+}$ samples.

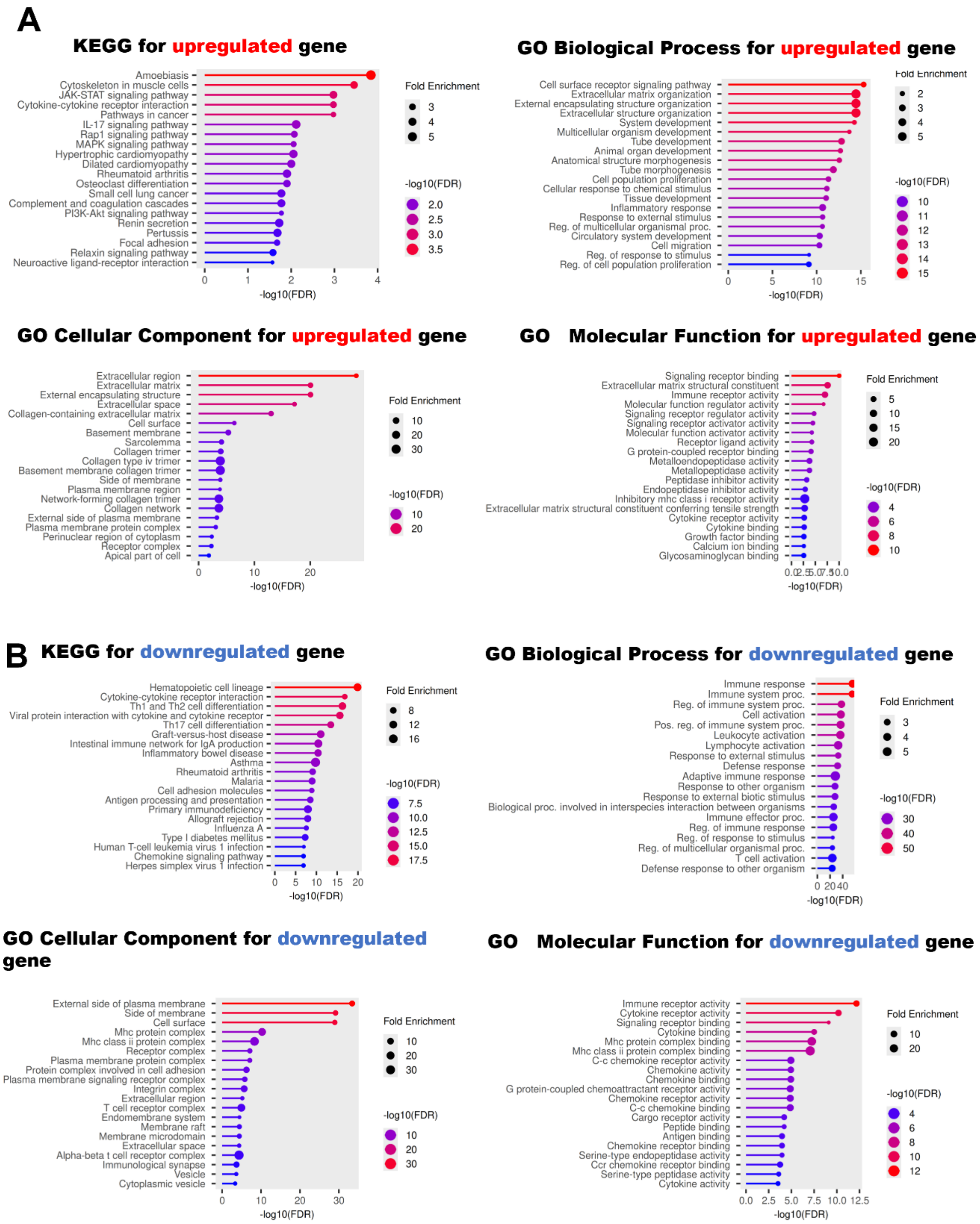


**Supplementary Figure 12. Functional enrichment analysis for testis immune cell transcriptome.**

(A) KEGG and GO analyses for upregulated genes showing enrichment in immune response pathways. (B) KEGG and GO analyses for downregulated genes.

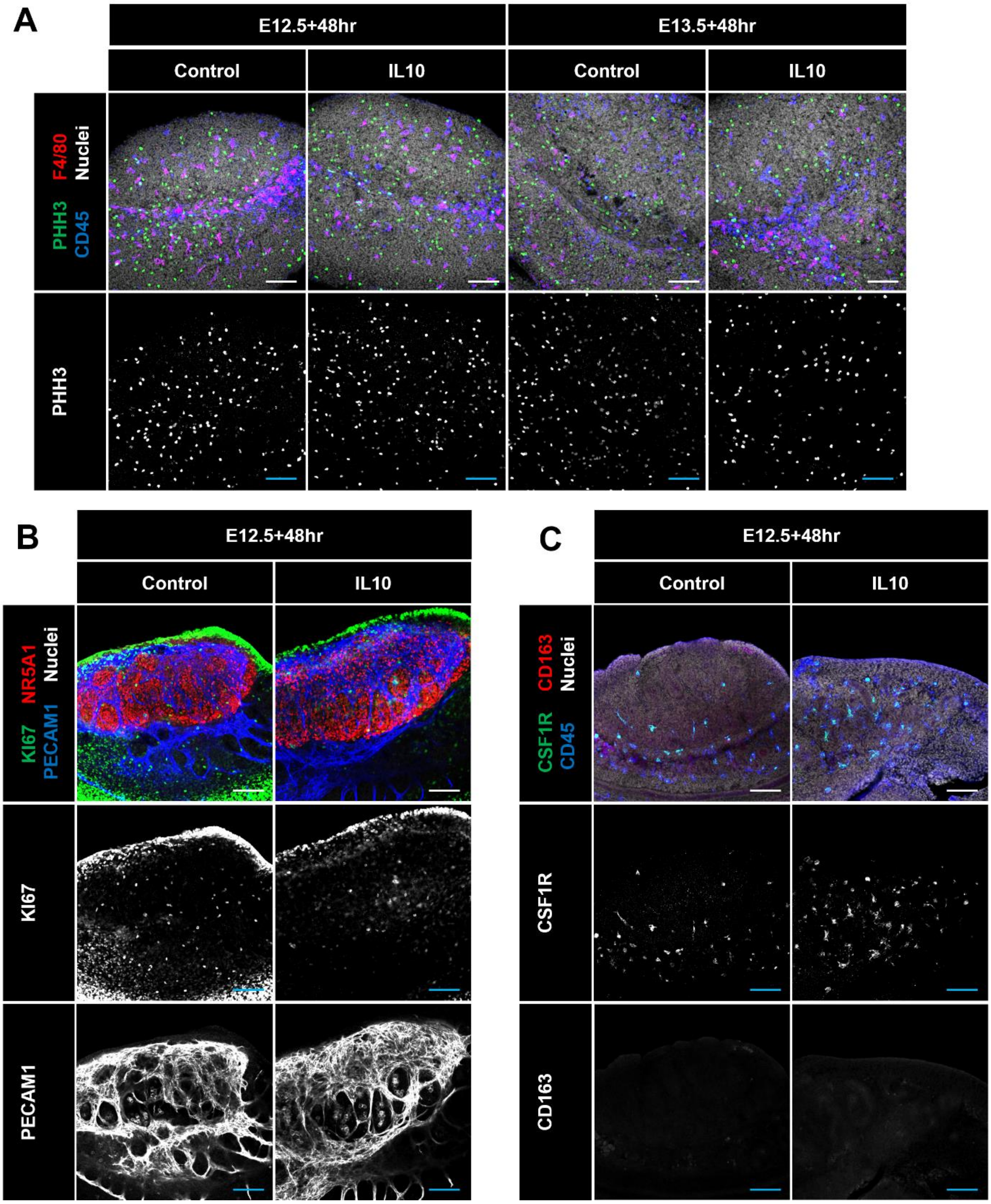


**Supplementary Figure 13. Effect of IL10 culture on immune and endothelial markers.**

(A) Immunostaining for PHH3/F4/80/CD45 in E12.5 and E13.5 testes cultured with IL10 for 48 hrs versus control testes. (B) Ki67/NR5A1/PECAM1 staining. (C) CSF1R/CD163/CD45 staining. Scale bar: 100 µm.

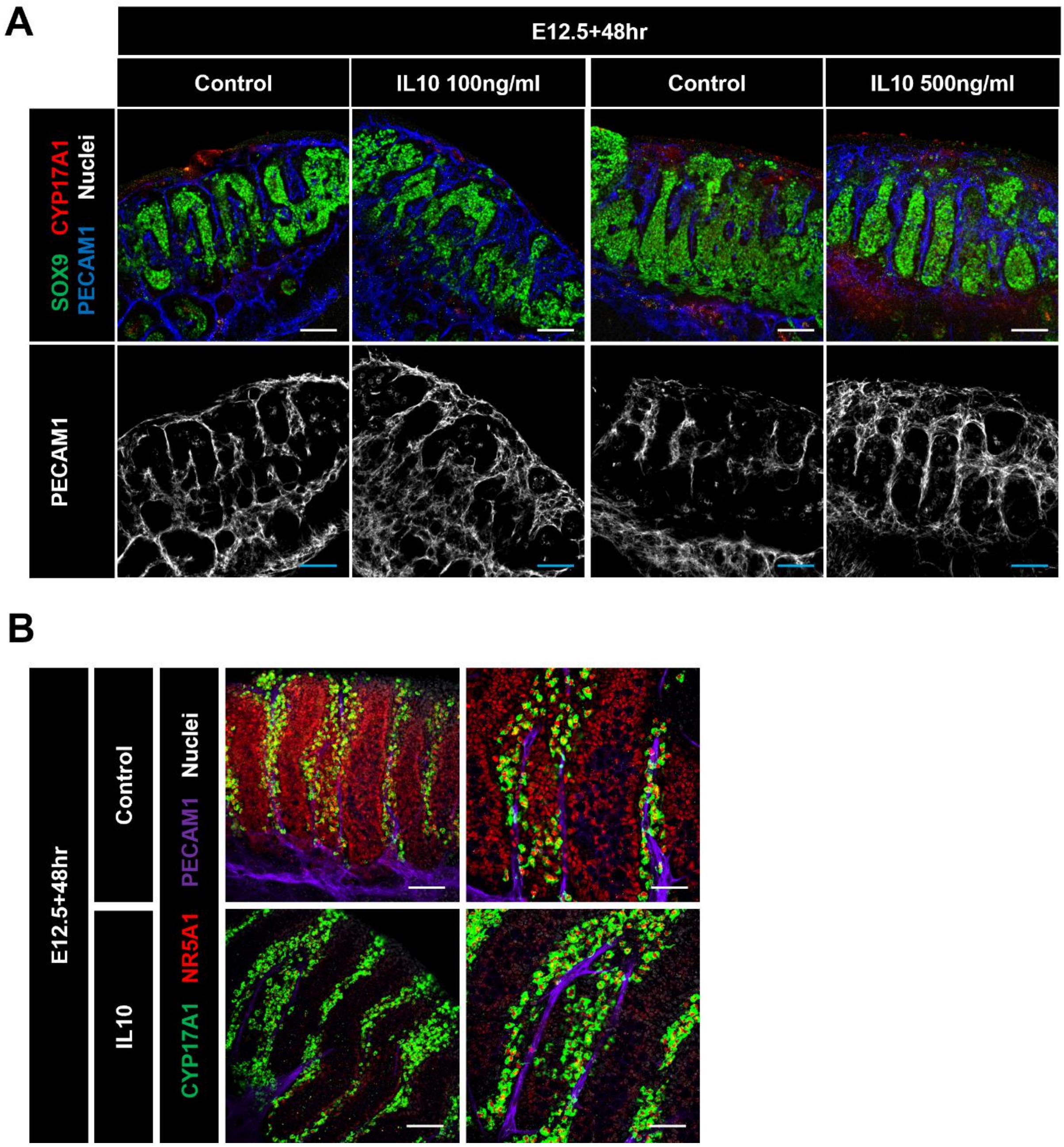


**Supplementary Figure 14. IL10 dose response on vascular and Leydig markers in cultured fetal testes.**

(A) Immunostaining for SOX9/CYP17A1/PECAM1 in E12.5 testes cultured with IL10 (100 or 500 ng/ml) for 48 hrs. (B) CYP17A1/NR5A1/PECAM1 staining. Scale bar: 100 µm.

**Supplementary Table S1. List of primary antibodies used for immunofluorescence.**

| Primary Antibody | Dilution | Source/Reference |
|---|---|---|
| Goat anti-JAG1 | 1:1,000 | R&D #AF599 |
| Mouse anti-NR2F2 | 1:500 | Perseus Proteomics #PP-H7147-00 |
| Rabbit anti-CYP11A1 | 1:2,000 | D. Wilhelm (1) |
| Rabbit anti-Nestin | 1:1,000 | BioLegend #PRB-315C |
| Rabbit anti-SOX9 | 1:500 | EMD Millipore #AB5535 |
| Rat anti-PECAM1 | 1:250 | BD Pharmingen #553370 |
| Goat anti-PECAM1 | 1:250 | R&D Systems #AF3628 |
| Goat anti-CYP17A1 | 1:500 | Santa Cruz #sc-46081 |
| Mouse anti-AMH | 1:500 | Santa Cruz #sc-365643 |
| Rabbit anti-pHH3 | 1:500 | Millipore #AF4465-SP |
| Rat anti-NR5A1 | 1:300 | Cosmo Bio #KAL-KO610 |
| Rabbit anti-StAR | 1:500 | Cell Signal Technology #8449 |
| Goat anti-CD45 | 1:500 | R&D #AF114 |
| Rat anti-CD45 | 1:300 | BioLegend #103101 |
| Rabbit anti-Cleaved Caspase 3 (Asp175) | 1:250 | Cell Signaling #9661S |
| Rabbit anti-CSF1R | 1:500 | Santa Cruz #sc-692 |
| Rabbit anti-ERG | 1:250 | Abcam #ab92513 |
| Rat anti-F4/80 | 1:2,000 | AbD Serotec #MCA497RT |
| Goat anti-FOXL2 | 1:250 | Novus #100-1277 |
| Rat anti-MKI67 (Ki67) | 1:500 | Thermo Fisher #14-5698-80 |
| Rat anti-TRA98 | 1:1,000 | Abcam #ab82527 |
| Goat anti-VCAM1 | 1:2,000 | R&D #AF643 |

**Supplementary Table S2. Sequences of primers used for qRT-PCR analyses.**

| **Gene name** | **Sequence (5' to 3')** |
|---|---|
| *Arx* forward | CAAGGATGGTGAGGACAGC |
| *Arx* reverse | TCTGGAACCACACCTGGACT |
| *Cdh5* forward | TCCTCTGCATCCTCACTATCACA |
| *Cdh5* reverse | GTAAGTGACCAACTGCTCGTGAAT |
| *Cyp11a1* forward | TGGCCCCATTTACAGGGAGAA |
| *Cyp11a1* reverse | GGCATCTGAACTCTTAAACAGGA |
| *Cyp17a1* forward | CAGAGAAGTGCTCGTGAAGAAG |
| *Cyp17a1* reverse | AGGAGCTACTACTATCCGCAAA |
| *Ddx4* forward | TACTGTCAGACGCTCAACAGGA |
| *Ddx4* reverse | ATTCAACGTGTGCTTGCCCT |
| *Gapdh* forward | AGGTCGGTGTGAACGGATTTG |
| *Gapdh* reverse | TGTAGACCATGTAGTTGAGGTCA |
| *Hsd3b1* forward | CAAGTGTGCCAGCCTTCATCT |
| *Hsd3b1* reverse | TTCATGATTCTGTTCCTCGTGG |
| *Insl3* forward | CCTCCTGGCTATGTCATTGC |
| *Insl3* reverse | CCTGTGGTCCTTGCTTACTG |
| *Jag1* forward | TGACATGGATAAACACCAGCA |
| *Jag1* reverse | GCAGCCCACTGTCTGCTATAC |
| *Lhx9* forward | TGTAATGCCCCAAGATTTGTTCTCCC |
| *Lhx9* reverse | ACCAGCAGCCTTATCCACCTTCACAG |
| *Nestin* forward | GCTGGAACAGAGATTGGAAGG |
| *Nestin* reverse | CCAGGATCTGAGCGATCTGAC |
| *Nr2f2* forward | GCCATAGTCCTGTTCACCTC |
| *Nr2f2* reverse | GCTCCTAACGTACTCTTCCAAAG |
| *Ren1* forward | CTCTCTGGGCACTCTTGTTGC |
| *Ren1* reverse | GGGAGGTAAGATTGGTCAAGGA |
| *Sox9* forward | GCGGAGCTCAGCAAGACTCTG |
| *Sox9* reverse | ATCGGGGTGGTCTTTCTTGTG |
| *Star* forward | TACATCCAGCAGGGAGAGGTG |
| *Star* reverse | CAGCGCACGCTCACGAAGTCT |
| *Adgre1 (F4/80) forward* | CCCCAGTGTCCTTACAGAGTG |
| *Adgre1 reverse* | GTGCCCAGAGTGGATGTCT |

| | |
|---|---|
| *Itgam (CD11b) forward* | CCACACTAGCATCAAGGGCA |
| *Itgam reverse* | AAGGGACACACTGACACCTG |
| *Ptprc (CD45) forward* | GGAGGACACAGCACATTGGA |
| *Ptprc reverse* | CCCCTGAGCAGCAATCATCA |
